# iFlpMosaics enable the multispectral barcoding and high-throughput comparative analysis of mutant and wildtype cells

**DOI:** 10.1101/2023.05.09.540000

**Authors:** Irene Garcia-Gonzalez, Stefano Gambera, Alvaro Regano, Susana F. Rocha, Lourdes Garcia-Ortega, Mariya Lytvyn, Luis Diago-Domingo, Maria S. Sanchez-Muñoz, Aroa Garcia-Cabero, Ivana Zagorac, Wen Luo, Macarena Fernández-Chacón, Verónica Casquero-Garcia, Federica Francesca Lunella, Rui Benedito

## Abstract

To understand gene function, it is necessary to compare cells carrying the mutated target gene with normal cells. In most biomedical studies, the cells being compared are in different mutant and control animals and therefore do not experience the same epigenetic changes and tissue microenvironment. The experimental induction of genetic mosaics is essential to determine a gene cell-autonomous function and to model the etiology of diseases caused by somatic mutations. Current technologies used to induce genetic mosaics in mice lack either accuracy, throughput or barcoding diversity. Here, we present a large set of new genetic tools and mouse lines that enable Flp recombinase-dependent ratiometric induction and single-cell clonal tracking of multiple fluorescently labeled wildtype and Cre-mutant cells within the same time window and tissue microenvironment. The labeled cells can be profiled by multispectral imaging or by FACS and scRNA-seq. This technology facilitates the induction and analysis of genetic mosaics in any cell type and for any given single or combination of floxed genes. iFlpMosaics enables a more accurate understanding of how induced genetic mutations affect the biology of single cells during tissue development, homeostasis, and disease.

## Introduction

The targeted deletion or mutation of a gene triggers a cascade of events in the cell that can significantly alter its phenotype and over time also the surrounding tissue. Scientists often analyze cells or tissues that have carried genetic mutations for several days, weeks, months, or even years and compare their phenotype with that of independent control cells or tissues from distinct non-mutant (control) animals. During the process from gene mutation to phenotypic manifestation and analysis, the biology of the targeted and surrounding tissue often undergoes significant changes that can interact with and alter the final phenotype of the mutant cells. An example of this is the conditional deletion or activation of genes important for vascular, neuronal, or cancer development. Mutations that impact tissue function or development frequently result in changes to the surrounding tissues. Since the non-targeted tissues that surround mutant cells are themselves a source of biochemical factors, any alteration to their development or function will trigger changes in a key tissue feedback mechanism; and any such changes will impact the phenotype of the mutant cells in a non-cell-autonomous manner that is independent of the initially induced genetic mutation itself (Hansen and Hippenmeyer, 2020; Hansen et al., 2022). Over time, this phenomenon often generates secondary mutant tissue phenotypes that can confound interpretation of the primary impact of a gene mutation on a cell’s phenotype.

Genetic mosaics are a powerful research tool for overcoming these problems because they allow the study of cell-autonomous genetic effects when mutant and control (wildtype) cells arise from the same progenitor cells and confront the same microenvironment in the same tissue or organism. In this scenario, the only difference between the cells being compared is the induced mutation, in an otherwise identical organism, genetic background, and tissue microenvironment.

Mouse models that allow the timed induction of somatic genetic mosaics are therefore essential for a comparative analysis of how mutant and control cell phenotypes develop over time in the same microenvironment. Such models also provide the only way to accurately study and model biological processes involving sporadic somatic mutations and their effect on cell proliferation, differentiation, or competition.

In *Drosophila*, interchromosomal mitotic recombination associated with distinct tissue markers has been widely used to induce and track genetic mosaics. A similar approach, called MADM (mosaic analysis with double markers), has been developed in mice to allow the labeling of control and mutant cells with different fluorescent markers (Contreras et al., 2021; Zong, 2014; Zong et al., 2005). This method allows full correlation between the expression of a given fluorescent marker and the genetic status (wildtype, heterozygous, or knockout) of the locus carrying the mutation, thus significantly decreasing the rates of false positives and false negatives. However, the method relies on difficult-to-achieve interchromosomal Cre-dependent recombination events, which overall leads to the generation of very few clones of labeled control and mutant cells in a Cre-expressing tissue. Due to the rarity of the interchromosomal recombination events, these genetic mosaics cannot be efficiently induced with tamoxifen-inducible CreERT2 lines, which are weaker and only transiently active. Moreover, this technology cannot be used in quiescent cells and requires chromosomal genetic linkage between the engineered MADM elements and another gene mutation, which significantly complicates the required mouse crosses and genetics, limiting its general applicability. MADM is also incompatible with epistasis analyses that require simultaneous loss-of-function of multiple genes present in distinct chromosomes.

Given the complexity and limitations of MADM, the most widely used method for generating conditional somatic genetic-mosaics in the mouse remains the much simpler CreERT2 recombinase-dependent mosaic induction of floxed-gene (flanked by loxP sites) deletion. With this method, the location, timing, and frequency of the recombination events can be regulated by restricting CreERT2 expression to a specific tissue and by varying the timing and dose of treatment with the CreERT2 activating ligand tamoxifen. This much simpler method requires the use of independent fluorescent reporters of Cre-recombination in order to detect cells with CreERT2 activity. However, several studies have shown that Cre-activity reporter alleles only accurately report recombination of themselves, and cannot be used to reliably report recombination of other floxed genes (Fernandez-Chacon et al., 2019; Liu et al., 2013; Schmidt-Supprian and Rajewsky, 2007). Thus, though much easier to implement than MADM, CreERT2-dependent genetic mosaic approaches generate a high frequency of false positives and false negatives and should not be used for mosaic genetics.

To overcome the limitations of current methods for the generation of genetic mosaics in mice, we have developed *iFlpMosaics*, a compendium of new genetic tools and mouse lines that enable FlpO-dependent-induction of multispectral and ratiometric genetic mosaics of both wildtype and mutant cells in the same tissue of an animal. *iFlpMosaics* provide high cellular barcoding and clonal resolution and tighter control of the frequency of genetic mosaicism over time. This technology is also compatible with all existing floxed alleles, and can be induced in any quiescent tissue or progenitor cells. The cell profiling and comparative analysis can be done by *in situ* multispectral imaging, or *ex situ* by FACS or single-cell RNA-seq (scRNAseq). *iFlpMosaics* will enable the accurate and high-throughput quantitative analysis how an individual or combinatorial set of genetic mutations affects the biology of single cells during tissue development, regeneration, or disease.

### False positives and false negatives with Cre-dependent mosaic genetics

CreERT2 genetics is prone to the induction of false positives and false negatives at high frequency. False positives are cells that recombine/express a reporter while failing to delete the independent floxed target reporter or gene. Conversely, false negatives are cells that delete the target floxed gene but do not recombine the specific reporter allele (Fig. 1a). The high frequency of false positive and false negative cells with CreERT2 genetics has been reported (Fernandez-Chacon *et al*., 2019; Liu *et al*., 2013), but so far no study has systematically quantified their occurrence for multiple reporter alleles and at the lower induction rates and tamoxifen doses required for mosaic genetics. In order to do this, we analyzed several independent mouse lines containing different Cre-reporters located in the Rosa26 locus, thus allowing us to detect false positives or false negatives among different Cre-reporters by FACS or histology. This analysis showed that different Cre-reporters recombine at significantly different relative frequencies, even when located in the same accessible Rosa26 locus and having similar genetic distances between loxP sites (Fig. 1b). As expected, the chance of finding cells with dual recombination of two reporters (true positives) decreased with decreasing tamoxifen dose and genetic mosaicism (Fig. 1c, 1d). Single-cell functional genetic or clonal analysis usually requires a decrease in the tamoxifen dose in order to ensure sparse labeling of mutant and wildtype cells in the tissue. Unequivocal identification of single cells and their clones over long periods of time requires a recombination rate of 0.01%-10%. We found that at these recombination frequencies the rate of true positives with CreERT2 genetics was very low (Fig. 1c, 1d).

**Figure 1:**
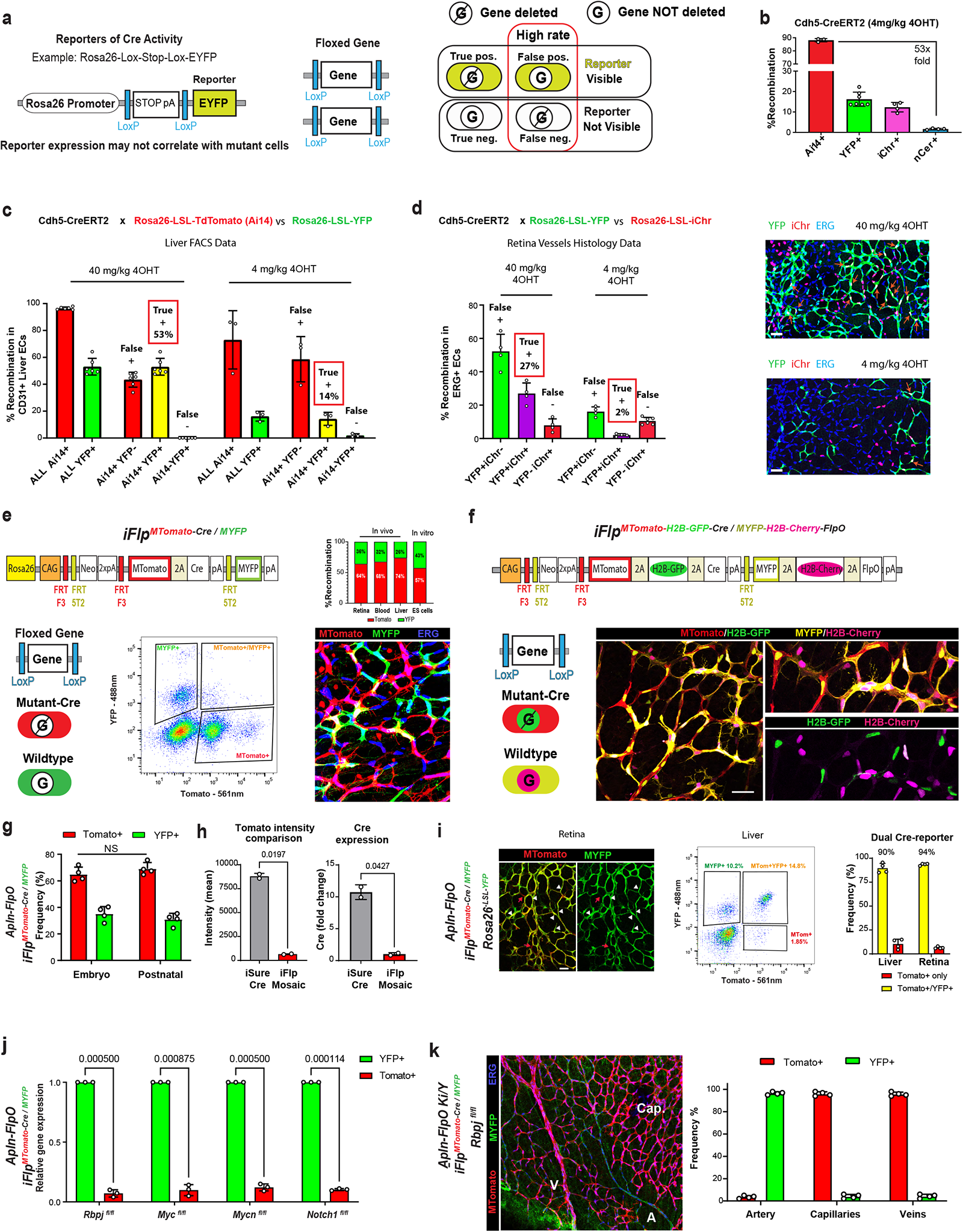
*iFlpMosaics* allow efficient and ratiometric labeling of mutant and wildtype cells. **a,** Schematic showing how a standard Cre-reporter can recombine and label cells without the deletion of any floxed gene (G) (false positives) and how the floxed gene can be deleted in non-Cre-recombined cells (false negatives). **b,** Different recombination efficiencies of Cre-reporters, despite all localizing to the *Rosa26* locus and having a similar genetic distance between LoxP sites. **c,d,** Cre-reporters only accurately report recombination of themselves, not that of other floxed alleles (reporters), particularly at low tamoxifen doses. **e,** The novel R26-*iFlp^MTomato-Cre/MYFP^* allele, in which Flpo/FlpOERT2-induced recombination of the mutually exclusive FRTF3 and FT5T2 sites leads to one of two possible outcomes: cells expressing MTomato and Cre (mutant) or cells expressing MYFP (wildtype) (see also Sup. Fig. 1). Induction leads to a ratiometric proportion of ∼ 65%MTom+:35%MYFP+. **f**, Tg-*iFlp^MTomato-H2B-GFP-Cre/MYFP-H2B-Cherry-FlpO^*allele and confocal micrographs showing the full spectral separation of the 4 fluorescent labels expressed in mutant and wildtype cells. **g,** In a wildtype background, the initial ratiometric proportion of MTomato+ and MYFP+ cells is maintained throughout embryogenesis to postnatal stages, indicating non-toxicity of Cre in MTomato+ cells. **h**, Comparison of MTomato intensity and Cre expression in cells of R26-*iFlp^MTomato-Cre/MYFP^* and Tg(*iSure-Cre*) mice. **i,** Efficient recombination of the Cre-reporter *Rosa26^-LSL-YFP^* in MTomato+ cells (white arrowheads) after induction of the R26-*iFlp^MTomato-Cre/MYFP^* allele by *Apln-FlpO*. There are few false positives (red arrows). The charts show representative FACS analysis of livers from the same animals and related numerical data from livers and retinas. **j,** Combination of the *iFlp^MTomato-Cre/MYFP^*allele with other floxed genes leads to very significant deletion of these in MTomato+ cells. **k,** Induction of *iFlp^MTomato-Cre/MYFP^* in mice carrying a floxed allele for *Rbpj*, an essential gene for arterial development, results in only 3.5% Tomato-2A-Cre+ cells in arteries (A), versus 96% in veins (v) and capillaries. Data are presented as mean values +/-SD. For statistics see Source Data File 1. Scale bars are 50μm.

These data show that most CreERT2-based *in vivo* functional genetic studies using mosaic genetics or sparse cell labeling will not achieve the desired targeted gene deletion in the labeled cells, and therefore will be unable to uncover the real effect of the targeted gene on single-cell biology with high accuracy and statistical power.

### Design and validation of *iFlpMosaic* mice

To overcome these problems with CreERT2-dependent mosaic genetic technologies, we developed a toolbox consisting of several mouse lines that allow the FlpO/FlpOERT2-recombinase-dependent induction of ratiometric mosaics of fluorescent mutant cells (gene KO) and normal cells (gene wildtype) (Fig. 1e, 1f and Sup. Fig. 1a, b). We call these new mouse alleles *iFlpMosaics* (*R26-iFlp^MTomato-Cre/MYFP^* and *Tg-iFlp^MTomato-H2B-GFP-Cre/MYFP-H2B-Cherry-FlpO^*). They contain the ubiquitously expressed CAG promoter and several distinct and mutually exclusive FRT sites (Cai et al., 2013) that enable multispectral labeling and fate-mapping of different mutant cells expressing the fluorescent membrane-tagged tomato protein (MTomato) and Cre, as well as of wildtype cells expressing membrane tagged YFP (MYFP), in any tissue expressing FlpO or FlpOERT2. The two *iFlpMosaic* lines differ either in the genomic location (ROSA26 versus random transgene) or in the expression of additional chromatin-bound fluorescent proteins (H2B-tag) for increased cell resolution and quantitative power (Fig. 1e, 1f and Sup. Fig. 1a, b).

Transfection of *iFlpMosaic* ES cells with FlpO-expressing plasmids or induction of these mosaics in mice carrying *FlpO/FlpOERT2* alleles induced the expected ratiometric mosaic of MTomato+ and MYFP+ cells, with very few cells being double positive due to recombination during the G2/M phase of the cycle (Fig. 1e, 1f, and Supplementary Fig. 1c-g). The observed higher frequency of MTomato+ cells is consistent with the shorter genetic distance between the FRTF3 sites than between the FRT5T2 sites (Fig. 1e and Sup. Fig. 1a, b, e). In ES cells and mice, the initially induced cell ratios did not change after prolonged culture or during tissue development *in vivo* (Fig. 1g and Sup. Fig. 1h), showing that expression of MTomato-2A-Cre (or MYFP) is not deleterious to cells. Notably, we found on average 10 times lower expression of the MTomato-2A-Cre cassette in *iFlpMosaic* mice than in the previously generated *iSuRe-Cre* mice (Fig. 1h), further reducing the possibility of Cre-toxicity (Fernandez-Chacon *et al*., 2019). This difference in expression is due to the different DNA expression elements and genomic location of these alleles.

In previous reports, inducible Cre-expressing alleles located in the Rosa26 locus were found to be leaky and to induce germline recombination, even in the absence of FlpO/CreERT2 activity (Fernandez-Chacon *et al*., 2019; Lao et al., 2012). Given that the *iFlpMosaic* alleles contain Cre-and FlpO-expressing cassettes that can be expressed if the upstream transcriptional stop signals are bypassed (as shown in previous mouse lines), we checked if *iFlpMosaic* mice were also leaky. None of the analysed *iFlpMosaic* mice had leaky expression of the fluorescent proteins contained in their constructs in the absence of an extra allele expressing FlpO (Sup. Fig. 1i). Crosses of *iFlpMosaic* mice with the *Rosa26^LSL-YFP^* Cre-reporter mice revealed a very low percentage of non-self Cre-leakiness in some organs of the *Tg-iFlp^Tomato-H2B-GFP-Cre/YFP-H2B-Cherry-FlpO^* mouse line, but not the *R26-iFlp^MTomato-Cre/MYFP^* line (Sup. Fig. 1i).

Significantly more important than false negatives is the occurrence of false positives. In *iFlpMosaic* mice, MTomato-2A-Cre+ cells efficiently recombined other Cre-reporter alleles and all floxed genes tested (Fig. 1i. 1j), ensuring high genetic reliability at single-cell resolution. For example, induction of *iFlpMosaics* in the *Rbpj* floxed background results in only 3.5% Tomato-2A-Cre+ cells in arteries versus 96% in veins and capillaries (Fig. 1k), indicating high gene-deletion efficiency of this master regulator of arterial development. Together, these data show that *iFlpMosaics* provide a very high rate of properly labeled and assigned mutant and wildtype cells. This is in stark contrast to standard CreERT2 mosaic genetics (Fig. 1c, 1d) and similar to the MADM system (Contreras *et al*., 2021; Zong, 2014; Zong *et al*., 2005). *iFlpMosaic* alleles thus enable the tightly controlled induction of reliable and ratiometric mosaics of mutant and wildtype cells in any tissue expressing FlpO or FlpOERT2; moreover, they provide both membrane and nuclear fluorescent markers for tracking mutant and wildtype cell proliferation, differentiation, or migration at very high resolution (Sup. Fig. 2).

### Generation of ubiquitous and tissue-specific FlpOERT2 mouse lines

Multispectral genetic mosaics are a particularly valuable research tool if they can be induced at a specific time point and in a significant number of cells, something that is not possible with the MADM system (Contreras *et al*., 2021; Zong, 2014; Zong *et al*., 2005). To obtain temporal control over Flp activity or induction, previous studies generated mouse lines that enabled the constitutive (Rosa26) or tissue-specific expression of FlpeERT or the mammalian codon-optimized FlpOERT (Hunter et al., 2005; Kranz et al., 2010). These studies showed that FlpOERT2 activity is significantly lower than CreERT2, and very few cells in each organ were shown to recombine (Lao *et al*., 2012). In agreement with this, we found that the published *R26^CAG-FlpOERT2^* line recombined the *R26-iFlp^MTomato-Cre/MYFP^* allele in only ∼0.2% of embryonic cells, and was also weak in postnatal and adult organs (Fig. 2a).

**Figure 2:**
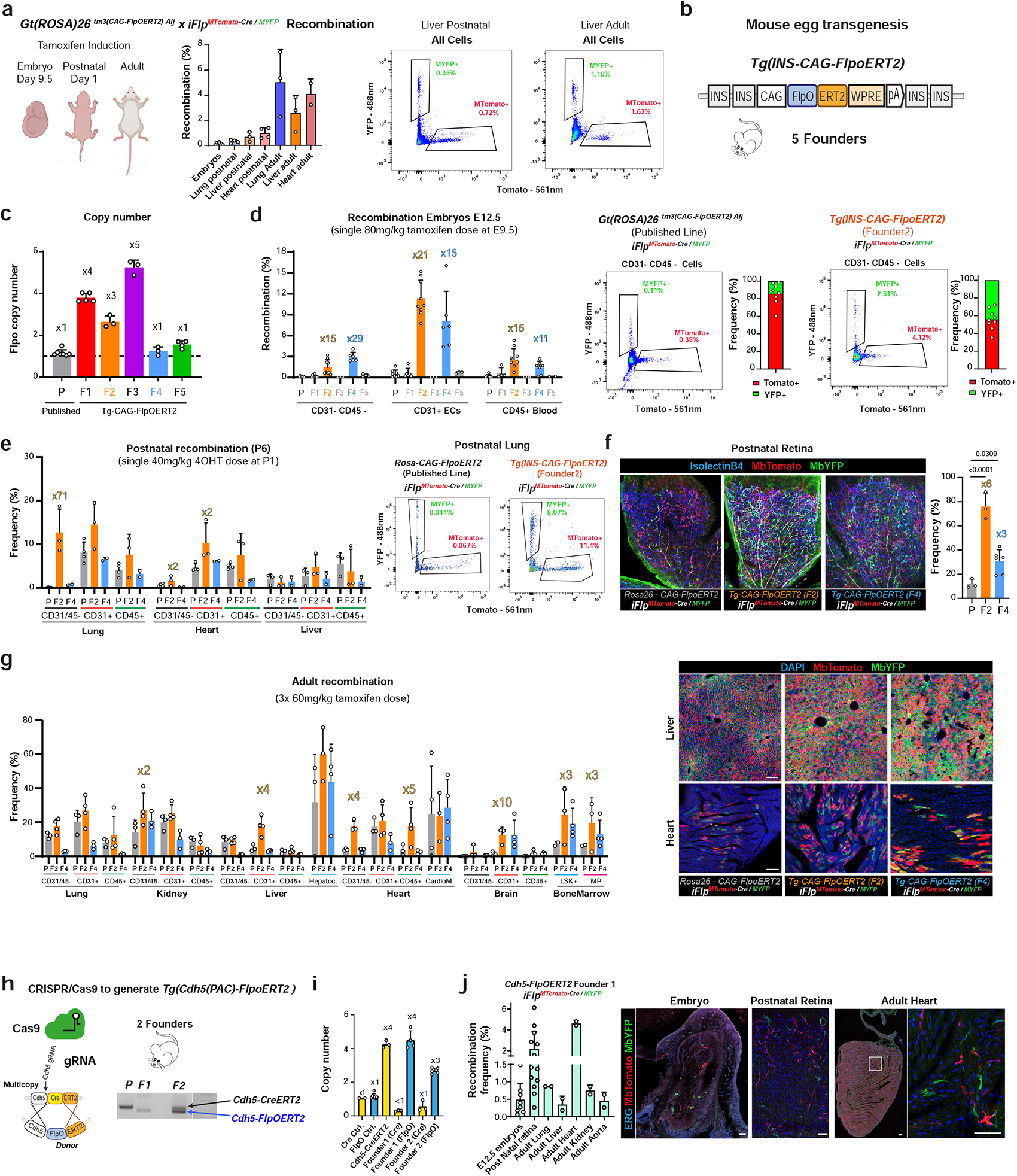
Generation of ubiquitous and tissue-specific FlpoERT2 mouse lines. **a,** Recombination of the *iFlp^MTomato-Cre/MYP^* mosaic mouse lines induced by the published *R26^CAG-FlpoERT2^* allele leads to reduced recombination efficiency at embryonic and postnatal stages and in adult organs. **b,** Schematic of novel *Tg(Ins-CAG-FlpoERT2*) alleles containing DNA elements optimized to enhance FlpOERT2 expression. **c,** Comparison of FlpO copy number by genomic DNA qRT-PCR. **d-g** Comparison by FACS and histology of the recombination efficiency of the different FlpOERT2-expressing mouse lines/founders at embryonic, postnatal, and adult stages. Fold change is indicated for the most significant changes. **h,** Crispr/Cas9 targeting procedure for the generation of novel endothelial-specific *Cdh5*-*FlpoERT2-* expressing mouse lines from the published *Cdh5-CreERT2* mouse line. **i,** Comparison of FlpO and Cre copy number by genomic DNA qRT-PCR. **j,** Recombination efficiency of Founder #1 of the newly generated *Tg(Cdh5-FlpoERT2)* mouse line in embryonic, postnatal, and adult tissues. Data are presented as mean values +/-SD. For statistics see Source Data File 1. Scale bars, 100μm.

To enhance FlpOERT2 expression, we therefore sought to develop new constructs containing optimized DNA elements (Cai *et al*., 2013; Pontes-Quero et al., 2017) (Fig. 2b). We generated and screened 5 transgenic ubiquitous-FlpOERT2 mouse lines with variable copy number (Fig. 2c). Compared with the established *R26^CAG-FlpOERT2^* line, the best of these new *Tg(Ins-CAG-FlpOERT2)* lines (here called *Tg(Ins-CAG-FlpOERT2)*^#F2^) induced several-fold higher recombination of the *iFlpMosaic* allele in embryos and different organs and cell types (Fig. 2d-g and Sup. Fig. 3a-c). With this higher recombination frequency, we frequently observed mutant (MTomato+) and wildtype (MYFP+) cells close to each other, enabling direct comparison of cells experiencing the same tissue microenvironment (Fig. 2f, 2g, Sup. Fig. 3d). As expected, the recombination rate differed between cell types and organs. We confirmed that the ubiquitous and strong *FlpOERT2* expression is not leaky (Sup. Fig. 3e).

Besides generating ubiquitous *FlpOERT2* lines of general relevance, we also developed a method to easily target and modify pre-existing and pre-validated *CreERT2*-expressing transgenic alleles to achieve tissue-specific expression of *FlpOERT2*. We designed unique guide RNAs and donor DNAs containing FlpO that allow the efficient conversion of any established *CreERT2* allele into a *FlpOERT2* expressing allele (Fig. 2h). Using this method, we converted an existing PAC *Cdh5-CreERT2* mouse line (Wang et al., 2010) containing 4 copies of the *CreERT2* sequence into PAC *Cdh5-FlpOERT2* mouse lines containing 3 or 4 copies of the *FlpOERT2* sequence (Fig. 2i). Of the two founder lines obtained, only the *Cdh5-FlpOERT2^F1^* line had no residual Cre sequence or activity (Fig. 2h). As expected, this line was expressed and induced specific recombination in endothelial cells (ECs, Fig.2j).

### Ratiometric functional genetics with *iFlpMosaic* mice

Unlike classical Cre-reporters or the *iSuRe-Cre* allele, *iFlpMosaics* are ratiometric by design. This allows quantification by histology or FACS of the relative frequencies of mutant (MTomato-2A-Cre+) and wildtype (MYFP+) cell populations induced in the same tissue and experiencing the same microenvironment throughout the pulse–chase period. We envisioned that this property would allow high-throughput and accurate analysis of the cell-autonomous function of genes in virtually all cell types and over long periods. Such an analysis is not possible with classical Cre/CreERT2 mosaic genetics, because of the high frequency of false positives and negatives (Fig. 1c, 1d) and because Cre-ERT2 genetics does not enable the simultaneous labelling of wildtype cells, which provide an important internal control.

To demonstrate the power of *iFlpMosaic* mice to reveal the effect of induced mutations in multiple cell lineages, we intercrossed them with mice containing either the *Tg(ACTFLPe)9205Dym* allele (Rodriguez et al., 2000) (abbreviated here as *ACTB:FlpE*; this allele recombines early embryo progenitor cells), the *Apln-FlpO* allele (Luo et al., 2021) (recombines endothelial and derived hematopoietic cells), or the new *Tg(Ins-CAG-FlpOERT2)* allele (induces recombination at any stage and in all cell types). These alleles were further combined with floxed alleles for the *Myc* and *Foxo* genes, which are considered to be important for the proliferation and differentiation of most cell types (Claveria et al., 2013; Eijkelenboom and Burgering, 2013).

When the *R26-iFlp^MTomato-Cre/MYFP^* allele was combined with the *ACTB:FlpE* or *Apln-FlpO* alleles and the *Myc and Foxo* floxed alleles, mosaic mutant animals survived until adult stages, unlike embryos carrying mutations of these genes in all cells or in all ECs (Dharaneeswaran et al., 2014; He et al., 2008). To understand how these genes affected the biology of different cell types, we first conducted a high-throughput FACS analysis to score the relative ratios of mutant cells (MTomato-Cre+) and wildtype cells (MYFP+) in major organs of postnatal day 7 animals. The assumption was that if the targeted gene is important for a given cell type or organ, its deletion would change the proliferation or differentiation rate of the mutant cells, and this would be noticed in the final MTomato+/MYFP+ log ratio detectable by FACS. This analysis revealed that the impact of MYC loss depended on the cell type and organ. MYC loss has previously been linked to a general competitive disadvantage (Claveria *et al*., 2013) and a lack of proliferation by ECs (Wilhelm et al., 2016). However, we found that mosaic *Myc* deletion had a relatively minor effect on EC clonal expansion throughout embryonic development (Fig. 3a). Instead, the highest sensitivity to the mosaic loss of MYC was observed in the hematopoietic lineage (Fig. 3b and Sup. Fig. 4a, b). Unlike liver and lung, heart CD31-CD45-cells were not significantly affected by *Myc* deletion (Fig. 3c), likely due to compensation by the homologous *Mycn* gene in cardiomyocytes as previously reported (Munoz-Martin et al., 2019).

**Figure 3:**
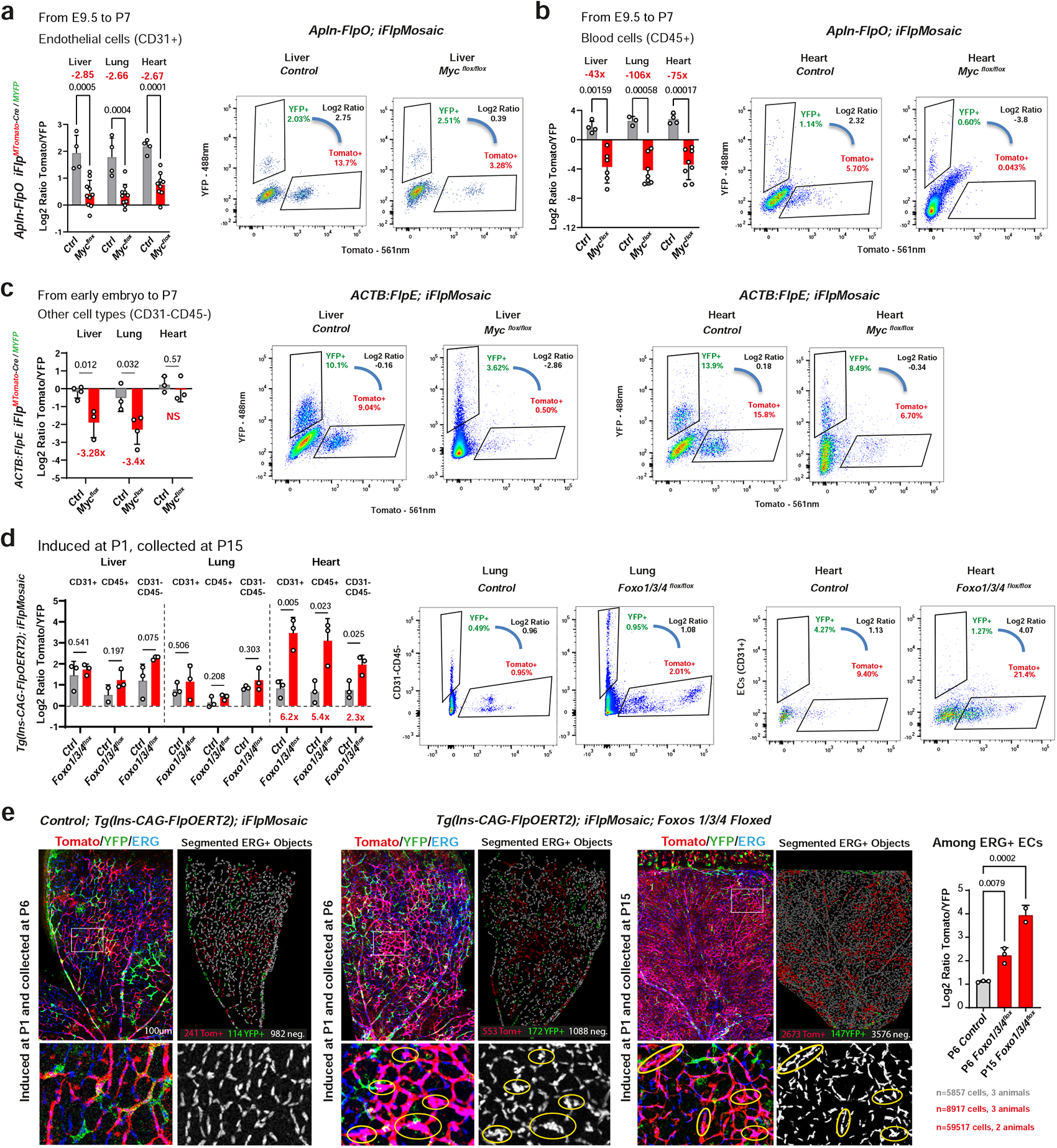
Ratiometric iFlpMosaics allow high-throughput assessment of a cell-autonomous gene function. **a,b**, *iFlp^MTomato-Cre/MYP^* mosaics were induced with *Apln-FlpO* (recombines endothelial and derived hematopoietic cells at E9.5) on control (wildtype) and Myc floxed backgrounds, and tissues were analyzed at P7. FACS data for CD31+ (ECs) and CD45+ (Blood) cells shows that *Myc* deletion (in MTomato+ cells) leads to a competitive disadvantage, especially in blood cells. Absolute fold change is indicated in red. **c,** When *iFlp^MTomato-Cre/MYFP^* mosaics are induced in most embryonic cells with *ACTB:FlpE*, analysis of other organ cell types (CD31-CD45-) reveals that *Myc* is necessary for the development of liver cells, but not lung or heart cells. **d,** *iFlp^MTomato-Cre/MYP^* mosaics were combined with *Tg(Ins-CAG-FlpOERT2)* on control (wildtype) and *Foxo1/3/4* floxed backgrounds, and tamoxifen was injected at P1 to induce recombination in all cell types. Liver, lung, and heart cells were analyzed at P15 by FACS. *Foxo1/3/4* deletion (in MTomato+ cells) increases the proliferation of heart cells, but not liver or lung cells. **e,** Representative confocal micrographs of control and *Foxo1/3/4* mosaic P6 and P15 retinas from animals induced at P1. The chart shows ratiometric analysis of retinal immunostainings for Tomato, YFP, and ERG (labels EC nuclei), showing that *Foxo1/3/4^KO^*ECs have a competitive advantage. Data are presented as mean values +/-SD. For statistics see Source Data File Scale bars are 100μm.

We also confirmed the ability of the system to support mosaic epistasis analysis of mutant cells having multiple deleted genes, in this case 3 *Foxo* genes (6 floxed alleles). Foxo genes partially compensate each other’s function in several cell types, and together they are considered important negative regulators of cell proliferation and metabolism downstream of AKT signaling, preventing uncontrolled cell growth (Eijkelenboom and Burgering, 2013; Paik et al., 2007; Wilhelm *et al*., 2016). Surprisingly, our analysis showed that mosaic deletion of three *Foxo* genes (*Foxo1/3/4^KO^*) had relatively minor consequences for cell expansion in the liver and lung, but especially affected heart cells (Fig. 3d). This suggests cell-type and organ-specific functions for *Foxo* genes. The data also show that postnatal retinal ECs are especially responsive to the loss of *Foxo1/3/4*. These cells were much more proliferative, formed dense clusters, and outcompeted wildtype cells over time (Fig. 3e), in line with the vascular phenotypes and hemangiomas shown previously using standard CreERT2 genetics (Paik *et al*., 2007; Wilhelm *et al*., 2016).

Overall, these results show the utility of *iFlpMosaics* technology for the accurate modeling and high-throughput analysis by FACS or tissue imaging of the impact of somatic mutations on cellular expansion and competition. In this particular case, *iFlpMosaics* analysis revealed that MYC and FOXO proteins are dispensable for the proliferation of many cell types in diverse organs, with MYC being more important in the blood lineage and FOXOs in retina ECs and heart cells.

### Combining *iFlpMosaic with iFlp^Chromatin^* for higher single-cell clonal resolution

An important potential application of *iFlpMosaics* technology is the direct live imaging of Cre-mutant and adjacent fluorescent wildtype cells. However, although live microscopy imaging is routine for many species and developmental model systems, it is still not possible or practical for most mouse tissues and laboratories due to the deep tissue location of most cells, the lack of appropriate microscopes, and the relatively slow pace of the biological processes under study. Analysis of how single cells proliferate, differentiate, and migrate in a mouse tissue requires advanced pulse–chase genetic and imaging tools. The *Brainbow* and *Confetti* genetic tools and *Dual ifgMosaics* were generated for this purpose (Livet et al., 2007; Pontes-Quero *et al*., 2017; Snippert et al., 2010). Compared with standard unicolor or single-molecule reporters of Cre activity, multispectral mosaics allow more precise and quantitative single-cell, retrospective, and clonal lineage tracing, due to the lower probability of a cell having a given stochastic recombination event or barcode. To increase the quantitative single-cell resolution of the *iFlpMosaics* technology, we combined the *Rosa26-iFlp^MTomato-Cre/MYFP^*allele with the published *Rosa26-iChr2-Mosaic* allele (also named here as *iCre^H2B-Cherry/GFP/Cerulean^*, see Sup. Fig. 4c) (Pontes-Quero *et al*., 2017). This enabled the stochastic and mutually exclusive expression of fluorescent proteins located in nuclei (for cell unit counts) or the cell membrane (for cell shape) in mutant cells (MTomato-2A-Cre+), expanding the diversity of cell barcoding from 1 to 3 possible mutant cells (Fig. 4a).

**Figure 4:**
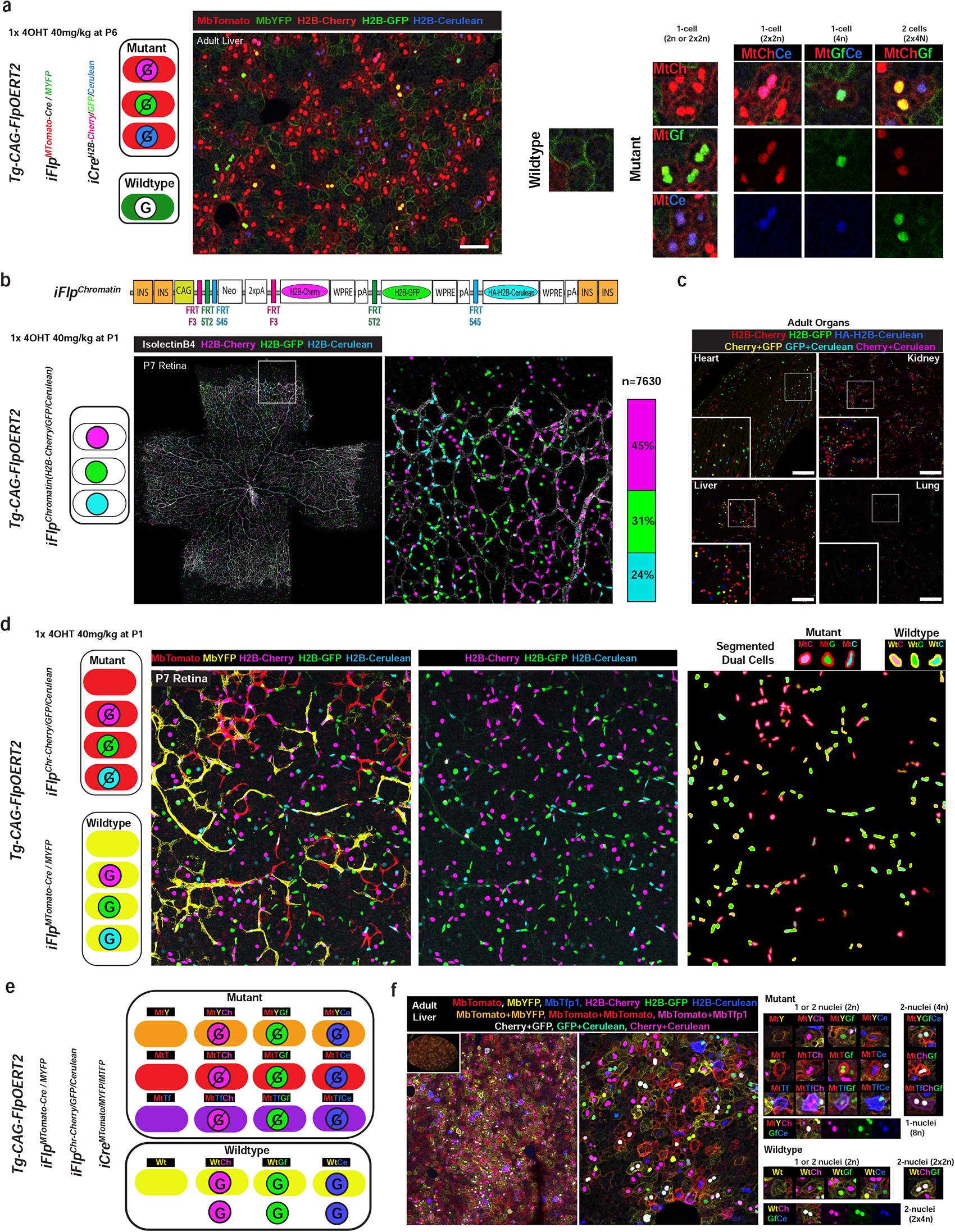
Combining *iFlpMosaic* with *iFlp^Chromatin^* yields higher single-cell clonal resolution. **a**, Representative confocal micrograph of adult liver sections from mice carrying the *Tg(Ins-CAG-FlpoERT2), iFlp^MTomato-Cre/MYFP^*, and *Rosa26-iChr2-Mosaic (iCre^H2B-Cherry/GFP/Cerulean^*) alleles. Combination of these membrane and nuclear reporter alleles allows up to 3 mutant and 1 wildtype cell barcodes (labeling). Note that binucleated (2x2n) or polyploid (4n) hepatocytes can have more multispectral barcodes. **b,** Representative confocal micrograph of a P7 retina from an animal containing the *Tg(Ins-CAG-FlpOERT2)* allele and the new *iFlp^Chromatin^* nuclear reporter allele (for explanations of similar DNA elements, see Sup. Fig.4c) induced at P1 with 4-OHT. The bar shows the ratiometric frequency of each nuclear/H2B+ fluorescent marker. **c,** Representative confocal micrograph of the indicated adult organ sections of animals carrying the *iFlp^Chromatin^* and *Tg(Ins-CAG-FlpOERT2)* alleles. Mononucleated diploid cells (kidney and lung) express only 1 of 3 possible nuclear/chromatin markers, whereas multinucleated or polyploid cells (hepatocytes and cardiomyocytes) can express a combination of 2 or 3 possible markers (6 possible chromatin barcodes in total). **d,** Recombination and cell barcoding possibilities in animals carrying the indicated alleles. The representative confocal micrographs of a P7 retina show that wildtype and mutant cells can both be labeled in 4 different ways, significantly increasing single-cell clonal resolution. The panel to the right illustrates how a Fiji image analysis script automatically detects, segments, and pseudocolors the nuclei of dual-labeled cells (nuclei and membrane). **e,** Recombination and diploid cell barcoding possibilities in animals carrying the indicated alleles. The representative confocal micrographs of an adult liver (collected 2 weeks after tamoxifen induction) show that mutant hepatocytes can be labeled with up to 16 different fluorescent barcodes, depending on whether the cells are mononucleated (diploid) or multinucleated (polyploid). Wildtype hepatocytes can have up to 8 different fluorescent barcodes (only 6 shown). Scale bars, 100μm.

Despite its higher clonal resolution for single MTomato+ mutant cells, this method does not permit quantification of the clonal expansion of single MYFP+ wildtype cells in the same microenvironment and at the same level of resolution. To obtain higher clonal resolution in wildtype cells, we developed the *iFlp^Chromatin^*mouse line (also called *iFlp^H2B-Cherry/GFP/Cerulean^*), in which three distinct and incompatible FRT sites enable the generation of a Flp-recombinase-dependent mosaic of cells expressing three distinct chromatin-localized (H2B-tag) fluorescent proteins (Fig. 4b, 4c). When crossed with the *Tg(Ins-CAG-FlpOERT2)* and *iFlp^MTomato-Cre/MYFP^-Mosaic* lines, this new *iFlp^Chromatin^* allele allowed us to generate up to 8 dual-labeled cells (4 mutant and 4 wildtype), significantly increasing the single-cell clonal lineage tracing resolution of mutant and wildtype cells in the same tissue microenvironment (Fig. 4d). These alleles can also be combined with the published Cre-dependent Tg(*iMb2-Mosaic*) allele (or *iCre^MYFP/Tomato/MTFP1^*) (Pontes-Quero *et al*., 2017) to provide even higher single-cell or clonal resolution of mutant cells (Tomato+ with 12 different fluorescent barcodes) and wildtype cells (YFP+ with 4 different fluorescent barcodes or YFP-with 3 different barcodes) (Fig. 4e). When compared with MADM, this technology provides significantly higher barcoding diversity and clonal resolution of both mutant and wildtype cells, and also allows the labeling and identification of binucleated and polyploid cells, such as adult liver hepatocytes (Fig. 4f).

### Single-cell functional mosaic genetics with *iFlpMosaics*

As an example, we show here how these new mouse alleles and technology enabled us to accurately quantify how single mutant and wildtype cells clonally expand, migrate, and survive when induced at the same time and in the same tissue microenvironment.

In the liver, a high percentage of neonatal hepatocytes are in S-phase (EdU+) or in cycle (Ki67+) at P1 and P7, and most stop dividing before P20 (Fig. 5a). However, hepatocyte single-cell clonal expansion dynamics are still unknown. *iFlpMosaic x iFlp^Chromatin^* mice were induced at P1, and livers were collected at P20 for multispectral imaging and single-cell clonal analysis. Multispectral confocal scanning and spatial mapping of clones revealed that hepatocyte proliferation (clones with 2 cells or more) was dispersed throughout the liver (Fig. 5b and Sup. Fig. 5a,b). The recombination frequency varied among animals, but the average clone size obtained was still similar (Fig. 5c), confirming that there is enough multispectral clonal and quantitative resolution at both low and high *iFlpMosaic* recombination frequencies. These data revealed that 17.2% of neonatal hepatocytes did not expand clonally, with most hepatocytes (69%) dividing only once or twice in 20 days. Only 2.4% of single hepatocytes yielded clones with 8 or more cells. The average clone size for wildtype cells was 2.9, in agreement with the observed 2.9-fold liver weight increase from P1 to 20 (Fig. 5c, d). Comparison of clonal expansion between wildtype (MYFP+) and mutant *Myc^KO^* (Mtomato-2A-Cre+) hepatocytes revealed that MYC is necessary for optimal postnatal hepatocyte clonal expansion. Mean *Myc^KO^*hepatocyte clone size was 2.25, and these clones generated only 125% more hepatocytes (125 new hepatocytes for every 100) over 20 days, whereas wildtype-cell clone size was 2.87, and these clones generated 187% more hepatocytes over the same period (Fig. 5e, f).

**Figure 5:**
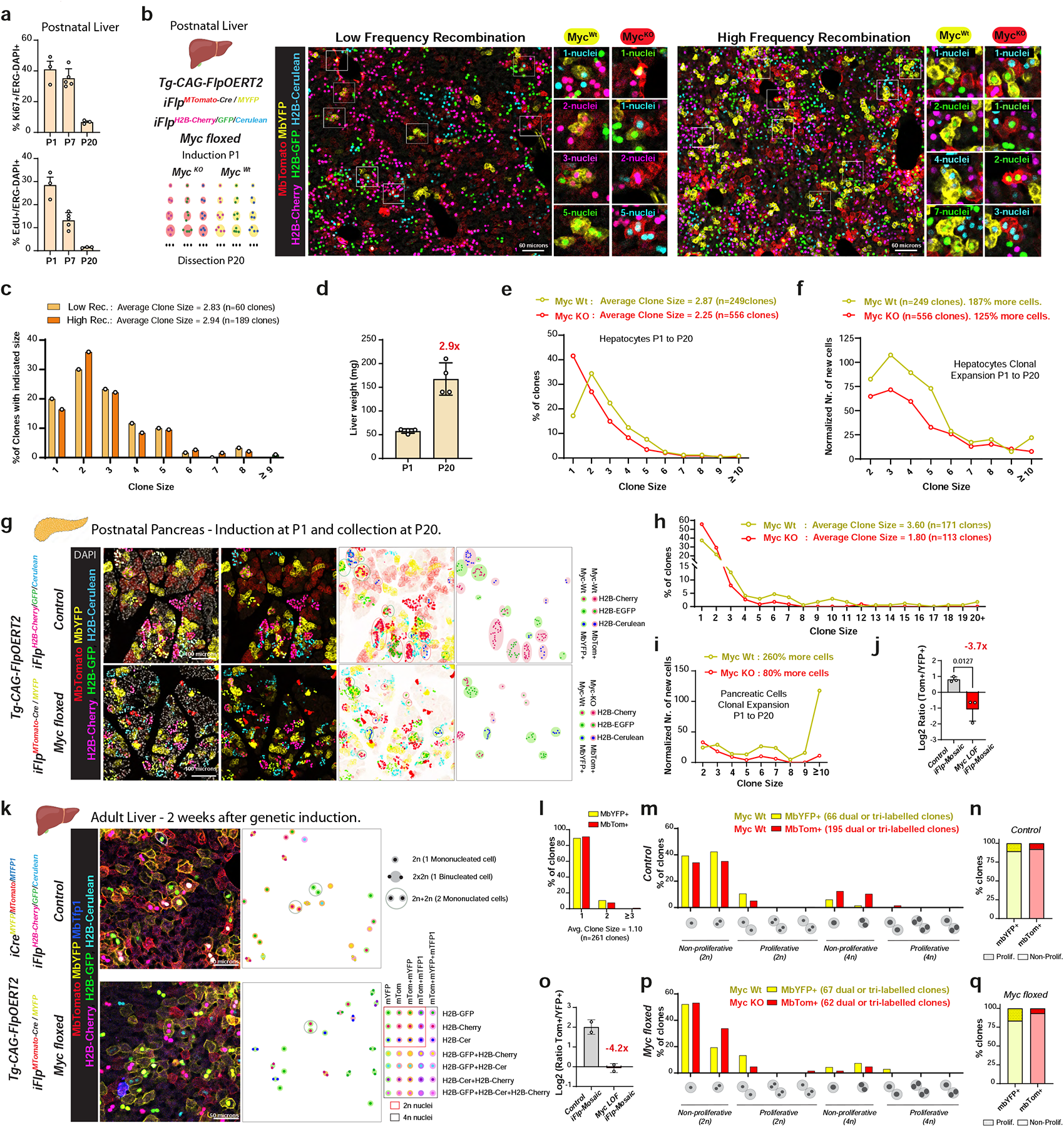
High-resolution *iFlpMosaics* reveal the impact of genetic mutations on single-cell clonal dynamics. **a**, Quantification of Ki67+ cells (in cycle) and EdU+ cells (S-phase) in P1, P7, and P20 livers. **b,** *Left* Schematic of the indicated alleles combined in a single animal and how they can be used to fluorescently barcode and track the clonal expansion of single mutant and wildtype cells from P1 to P20. *Right* Representative confocal micrographs of P20 livers (from animals induced with 4-OHT at P1) with different recombination frequencies (see also Sup. Fig. 5). **c,** Clone size distributions at both low and high recombination rates. Recombination frequency and fluorescent barcoding density do not affect clonal resolution and quantification. **d,** Liver weight at P1 and P20. **e,** Quantification of MYFP+ (*Myc^WT^*) and MTomato+ (*Myc^KO^*) clone size frequency, showing that *Myc^KO^*hepatocytes divide less. **f,** Chart depicting the normalized expansion of *Myc^WT^* and *Myc^KO^* hepatocytes over the same 20-day period. **g,** Representative confocal micrographs of P20 pancreas from control and *Myc*-floxed mice carrying the indicated *iFlpMosaic* alleles. Right panels are inverted LUT images depicting the clone mapping (Fiji image analysis scripts) according to their dual color code. **h,** Quantification of MYFP+ (*Myc^WT^*) and MTomato+ (*Myc^KO^*) clone size frequency, showing that *Myc^KO^* pancreatic cells divide less. **i,** Chart depicting the normalized expansion of *Myc^WT^*and *Myc^KO^* pancreatic cells over the same 20-day period. **j,** Log2 ratio of Tom+ cells to YFP+ cells in control and *Myc*-floxed pancreas, showing a 3.7 fold loss of MTomato+ (*Myc^KO^*) cells over 20 days. **k,** Representative confocal micrographs of adult livers from control and *Myc*-floxed mice in which the indicated *iFlpMosaic* and *iCre^Mb2-Mosaic^* alleles were induced 14 days before tissue collection. Right panels show automatic mapping of clones (Fiji image analysis scripts) according to their fluorescent barcode. Note the loss of MTomato+ cells in the *Myc*-floxed background. **l,** Clone size, showing that wildtype hepatocytes rarely divide or expand over a 2-week period. **m,** Clone frequency of wildtype MYFP+ and MTomato+ hepatocyte clones, showing the lack of impact of Cre expression in MTomato+ cells. Note the difference between diploid, mononucleated, polynucleated, and polyploid cells. **n,** Percentages of proliferative (composed of more than 1 cell) and non-proliferative YFP+ and Tomato+ clones, showing no major difference between the two wildtype populations. **o,** Log2 ratio of the Tom+ and YFP+ cell frequency in control and adult *Myc*-floxed livers, showing loss of a large fraction of *Myc^KO^* cells over 2 weeks. **p,** Clone frequency of YFP+ (*Myc^WT^*) and Tomato+ (*Myc^KO^*) hepatocytes, showing that the surviving *Myc^KO^*population contains an increased frequency of non-proliferating binucleated cells and a decrease frequency of proliferating mononucleated cells. **q,** Percentages of proliferative and non-proliferative YFP+ (*Myc^WT^*) and Tomato+ (*Myc^KO^*) clones, showing that loss of *Myc* reduces proliferative-clone frequency.

Single-cell clonal expansion analysis of postnatal pancreatic cells revealed similar Ki67+ and EdU+ frequencies to those of liver cells (compare Fig.5a with Sup. Fig. 5c, d). However, *iFlpMosaics* analysis revealed that wildtype pancreatic cells expand significantly more than hepatocytes until P20 (Fig. 5g), with a mean clone size of 3.60 instead of 2.87 (compare wildtype values in Fig. 5h,i with Fig.5e,f). As in the liver, *Myc^KO^* cells in the pancreas also expanded significantly less than wildtype cells (Fig. 5h, 5i) and were 3.7-fold less frequent than neighboring wildtype cells at the end of the analysis (Fig. 5j).

Evidence from recent studies suggested the existence of a highly proliferative hepatocyte population in adult liver, based on a new Dre-inducible Ki67-Cre (ProTracer) allele (He et al., 2021) or CreERT2-based single-cell lineage tracing of a unicolor reporter (Wei et al., 2021). Ki67 immunostaining showed that only 1.87% of adult liver hepatocytes are in cycle at any given moment (Sup. Fig.5d). Ki67 can be expressed in metabolically activated (primed and in cycle) or DNA-replicating (S-phase) or in polyploid cells, but many Ki67+ cells will not undergo productive cell division or will die after being in cycle. The *iFlpMosaics* system can measure both productive cell expansion and cell survival. Using the *iFlpMosaic* mice with the highest number of multispectral combinations, and therefore the highest single-cell clonal resolution, we observed very limited proliferation or clonal expansion of single mononucleated or binucleated hepatocytes during the 2 weeks after induction. The mean size of adult hepatocyte clones was 1.10 cells after 2 weeks (Fig. 5k, l), significantly lower than the 1.38 cells after only 1 week of pulsing reported for a simpler unicolor reporter (Wei *et al*., 2021). During the 2 weeks after induction, only 9.5% of single mononucleated or binucleated diploid hepatocytes underwent productive division or cytokinesis from binucleated cells to form a 2-cell clone (Fig. 5m, n). Proliferation frequency was also low in polyploid hepatocytes (4.9%).

Despite the absence of significant hepatocyte proliferation in homeostasis, when we induced *iFlp* ratiometric mosaics in adult *Myc^flox^* livers, we still observed a very significant loss of mutant *Myc^KO^* cells (4.2-fold decrease) only 2 weeks after induction (Fig. 5o). Since there is limited clonal expansion of wildtype cells in adult liver (mean clone size 1.10, meaning that only 10 new hepatocytes were generated for every 100 initial hepatocytes over 2 weeks; Fig. 5l), these data suggest that Myc loss induces a significant decrease in adult hepatocyte survival. Interestingly, wildtype hepatocytes expanded more when surrounded by *Myc^KO^*hepatocytes than when surrounded by wildtype hepatocytes (compare Fig. 5m/n with 5p/q and Sup. Fig.5e), probably to compensate for the loss of *Myc^KO^* hepatocytes. These data suggest the existence of Myc-dependent cell competition among adult hepatocytes, in which cells with lower Myc are less fit and are gradually excluded, as previously shown to occur during early embryonic development (Claveria *et al*., 2013). This competition and cell turnover may explain why liver cell mass does not increase over time, despite constant cell proliferation. Among the surviving *Myc^KO^* mutant cells, we also observed a significant increase in the percentage of binucleated cells and a decrease in the percentage of 2-cell mononucleated clones, suggesting that binucleated *Myc^KO^* hepatocytes undergo cytokinesis less frequently than their wildtype counterparts (Sup. Fig. 5e).

### *iFlp/Dre^Progenitor^* enables induction of genetic mosaics from single progenitor cells

The results above also show that there is significant intercellular clonal variability when different progenitor cells, occupying different tissue locations, are induced. This decreases the statistical robustness of gene-function analysis in single cells and requires a comparative analysis of a large number of single-cell derived clones and averaging of clone-size differences. With the above method, all Tomato+ and all YFP+ clonal populations in a given region can be averaged and compared. However, since the clones derive from different progenitor cells, is not possible to know exactly how loss of a gene changes the cellular and clonal expansion phenotype of a specific single-cell progeny located in a particular niche. The MADM approach allows the generation of labeled wildtype and mutant cells from the same progenitor cell, and in this way gives a very precise estimate of how a gene mutation impacts the mobilization and proliferation phenotypes of a single-cell derived progeny. However, as mentioned above, MADM is a cumbersome method, cannot be effectively induced at a specific time point, and is incompatible with the numerous existing floxed genes.

To overcome the limitations of current approaches to understanding the role of genes in single-progenitor cell biology, we designed the *iFlp/Dre^Progenitor^* DNA construct (Fig. 6a). Injection of this construct into mouse eggs produced four founder transgenic animals, only one of which worked properly and which we called *iFlp/Dre^ProgenitorF1^*. These *iFlp/Dre^Progenitor^* alleles can be induced by FlpOERT2 or the stronger DreERT2. Therefore, we also generated a new *Tg(Ins-CAG-DreERT2)*^#F4^ allele by using CRISPR/Cas9 assisted targeting into the pre-existing and screened *Tg(Ins-CAG-FlpOERT2)* allele (Fig. 6b). We reasoned that combination of these alleles with *iFlpMosaics* alleles would allow tamoxifen induction of a genetic cascade culminating in the generation of genetic mosaics of mutant and wildtype cells derived from the same progenitor cells. After tamoxifen induction of FlpOERT2 or DreERT2, the *iFlp/Dre^Progenitor^* allele is recombined, and this triggers the constitutive expression of FlpO, H2B-V5, and the cell-surface marker hCD2. FlpO then effectively recombines the *R26-iFlp^MTomato-Cre/MYFP^*allele in H2B-V5+/hCD2+ cells (Fig. 6c). We reasoned that since FlpO is a relatively weak recombinase and full transgene expression takes some time, there would be a corresponding delay in the recombination of the *R26-iFlp^MTomato-Cre/MYFP^*allele. During this period, a dividing cell will often generate two or more daughter cells, and in these, the *iFlpMosaic* allele can be recombined in one of two possible ways, generating either Tomato-2A-Cre+ or YFP+ cells in the progeny of a single cell.

**Figure 6:**
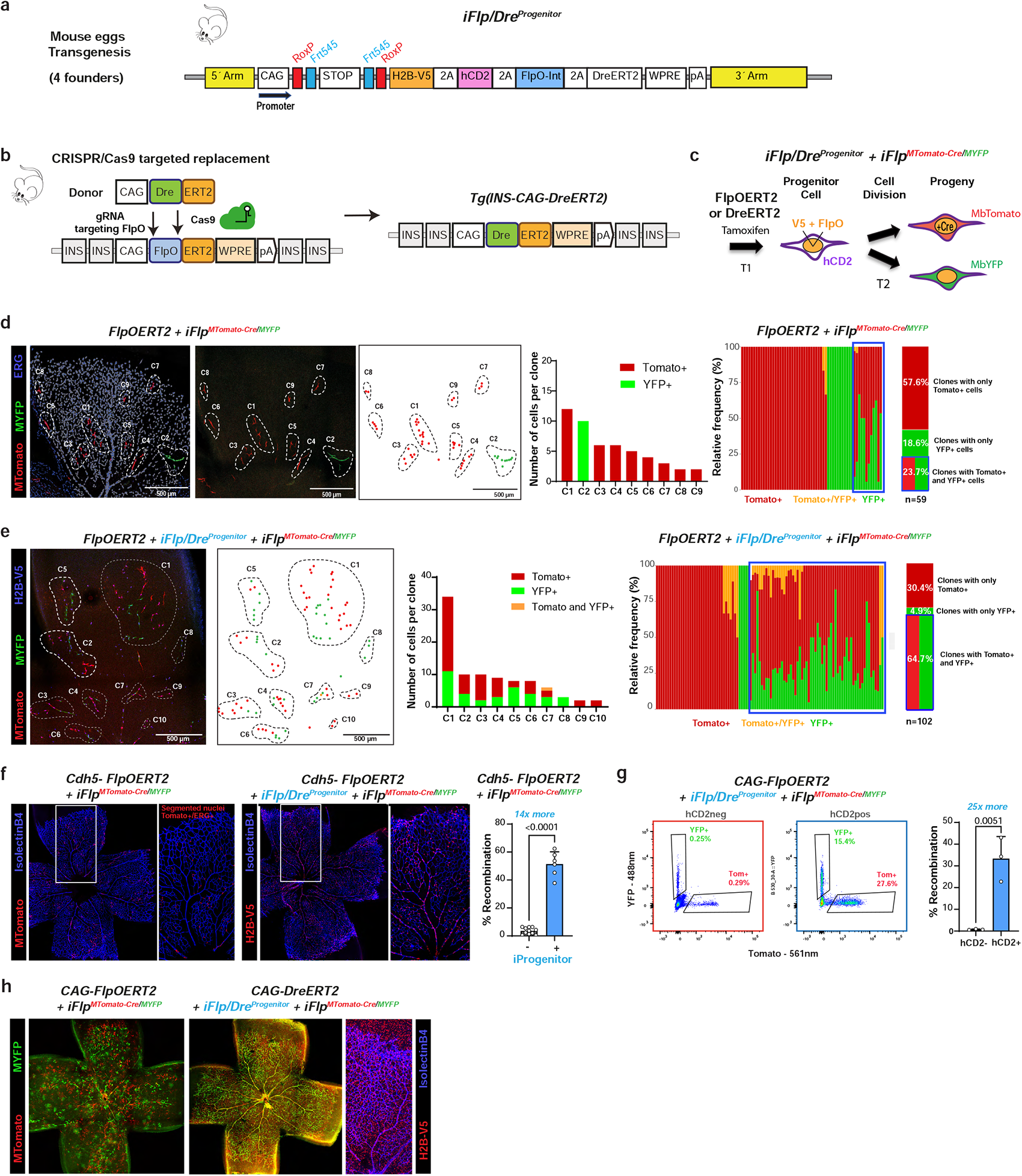
*iFlp/Dre^Progenitor^*enables the induction of genetic mosaics in the progeny of single cells. **a,** Schematic of the *iFlp/Dre^Progenitor^* DNA construct used to generate transgenic mice. **b,** Generation of *Tg(Ins-CAG-DreERT2)* by CRISPR-Cas9 mediated replacement of FlpO by Dre in the existing *Tg(Ins-CAG-FlpOERT2)^F4^*allele. **c**, Schematic of the genetic cascade induced in animals containing the indicated alleles. After tamoxifen induction of FlpOERT2 or DreERT2, the *iFlp/Dre^Progenitor^* allele is recombined, driving expression of the nuclear marker H2B-V5, the surface marker truncated human CD2 (hCD2), and FlpO. FlpO is a relatively weak recombinase that needs to be expressed and accumulate in the cell before it recombines the *iFlp^MTomato-Cre/MYFP^*allele. This delay results in recombination in the progeny of a single progenitor cell. **d**,**e**, In animals lacking the *iFlp/Dre^Progenitor^*allele, most single-cell derived clones are formed by just one type of cell (MTomato+ or MYFP+). With the *iFlp/Dre^Progenitor^* allele, it is possible to induce clones formed by mutant (MTomato+) and wildtype (MYFP+) cells derived from single progenitor cells (see also Sup. Fig. 6). **f**, Retinal confocal micrographs showing that the *iFlp/Dre^Progenitor^*allele significantly increases the recombination frequency of *iFlp^MTomato-Cre/MYFP^*(induced at P3 and collected at P7), enabling high frequency genetic mosaics with the weaker *Cdh5-FlpOERT2* line (expressed only in Cdh5+ ECs). H2B-V5 labels the nuclei of cells recombining and expressing the *iFlp/Dre^Progenitor^* allele. **g**, FACS plots showing that the *iFlp/Dre^Progenitor^* allele significantly increases the recombination frequency of the *iFlp^MTomato-Cre/MYFP^*allele when combined with the *Tg(Ins-CAG-FlpOERT2)* allele. **h**, Retinal confocal micrographs showing that combination of the new *Tg(Ins-CAG-DreERT2)* line with the *iFlp/Dre^Progenitor^* allele increases the recombination fequency of the *iFlp^MTomato-Cre/MYFP^* allele. H2B-V5 labels the nuclei of cells recombining and expressing the *iFlp/Dre^Progenitor^*allele.

To test this hypothesis, we injected tamoxifen into P4 mice and collected tissues for analysis 3 days later at P7. In animals without the *iFlp/Dre^Progenitor^* allele, Tomato+ and YFP+ cells arose from independent progenitors that were genetically pulsed in independent locations, and YFP+ wildtype cells were rarely intermixed with mutant Tomato+ cells (Fig. 6d). In animals with the *iFlp/Dre^Progenitor^* allele, anti-V5 immunostaining enabled the detection of the nuclei of the progenitor cell and its progeny. V5+Tomato+ and V5+YFP+ cells were frequently located close to each other, confirming that they arose from a single progenitor cell (Fig. 6e and Sup. Fig. 6a). This allows for direct comparisons of gene function in the progeny of single progenitor cells.

The *iFlp/Dre^Progenitor^* allele also allowed us to increase the sensitivity to FlpOERT2 recombination by 14-fold. With the Cdh5-FlpOERT2 line, we increased EC-recombination efficiency from 3.6% to 51% (Fig. 6f). Recombination efficiency was further increased by combining the *iFlp/Dre^Progenitor^* allele with *Tg(Ins-CAG-DreERT2)*^#F4^ instead of *Tg(Ins-CAG-FlpOERT2)*^#F4^ (Fig. 6g).

Another advantage of the *iFlp/Dre^Progenitor^* allele is the possibility of using anti-hCD2 magnetic beads or antibodies to rapidly isolate the recombined cells, even when they occur at low frequency. The hCD2+ cell population included a 25-fold higher percentage of Tomato+ and YFP+ cells (Fig. 6h), resulting in much faster isolation of these cells, an important consideration when many tissues or infrequent populations need to be isolated on the same day for scRNA-seq analysis.

### scRNA-seq of ratiometric *iFlpMosaics* to uncover gene function in all cell types

Recent advances in scRNA-seq methods have allowed unprecedented advances in the understanding of single-cell biology. However, there are three major limitations to the use of scRNA-seq technologies to assess the transcriptome of single mutant and wildtype cells resulting from conditional genetic experiments. The first limitation is the introduction of bias due to the profiling of cells from distinct mutant and wildtype animals that experienced different epigenetic and microenvironmental changes during their lifetime or in disease. The second is the reliance on standard Cre-reporters to define and isolate the mutant cells of interest, since these reporters do not correlate with the desired genetic deletions, particularly in CreERT2-inducible experiments (Fig. 1). The third and most important limitation is uncertainty as to whether the desired floxed gene will be deleted in the pool of analyzed and sequenced mutant single cells. This is important because scRNA-seq technologies only detect the mRNA of some genes in most single cells, and also because many floxed genes conserve their 3’ mRNA sequences after genetic deletion, leading to the transcriptional profile of pseudomutant cells whose true genetic status cannot be accurately determined by standard scRNAseq technologies.

Our analysis provides evidence that, unlike other genetic technologies, *iFlpMosaic* alleles allow the reliable isolation of *bona fide* mutant cells and wildtype cells from the same tissue (Fig. 1 and 2). Therefore, we combined *iFlpMosaics* with scRNA-seq to achieve high-throughput scoring of gene function in the proliferation and differentiation of all early embryo cell types. For this analysis, we selected to induce the deletion of *Rbpj*, which encodes a transcription factor essential for Notch signaling, one of the most important pathways for cell proliferation and differentiation during embryonic development (Siebel and Lendahl, 2017). We pulsed embryos with 4-OHT at embryonic day 9.5 (E9.5) and collected Tomato+ (mutant) and YFP+ (wildtype) cells by FACS at E13.5 (Fig. 7a). Based on their transcriptome and known marker genes, we identified 16 major cell types (Fig. 7b). In Tomato-2A-Cre+ cells, deletion of *Rbpj* exons 6 and 7 generates a less stable but still detectable 3’mRNA (Fernandez-Chacon et al., 2023), together with a decrease in the expression of the main canonical downstream target *Hes1* (Fig. 7c). The canonical *Hes1* target gene can also be regulated by other pathways. Deletion of *Rbpj* led to changes in the frequency of cells in S-phase (Ki67+ mRNA) in some clusters (Fig. 7c). Global *iFlpMosaic-*driven *Rbpj* deletion at E9.5 compromised the differentiation and expansion of peripheral neurons, ECs, cardiomyocytes, erythrocytes, hepatocytes, and myocytes, whereas the frequency of epithelial and mesenchymal cells increased (Fig. 7d-e). *Rbpj* deletion led to a 5-fold decrease in the frequency of ECs (Fig.7d, e), loss of expression of the arterial marker genes *Gja5* and *Gja4*, and upregulation of the tip-cell genes *Esm1, Kcne3, Apln*, and *Cdkn1a* (Fig.7g,h). This correlates with the essential role of *Rbpj* in the specification of arterial ECs and the inhibition of tip ECs (Luo *et al*., 2021; Pontes-Quero et al., 2019). *Rbpj^KO^* ECs also upregulated genes related to epithelial-to-mesenchymal transition, glycolysis, and angiogenesis and were less frequently in cycle (Fig.7i). scRNAseq data from other cell types revealed that *Rbpj* loss changes the relative frequency of specific cell-type clusters, suggesting an impact on their proliferation or differentiation. Indeed, *Rbpj* deletion dysregulated multiple genes and genetic pathways in the many cell types and clusters identified (Fig. 7j-r and Sup. Fig. 7). These scRNA-seq data provide an important resource for further exploration of the role of Rbpj–Notch signaling in the early expansion and differentiation of embryonic lineages. This analysis also exemplifies how the combination of ratiometric *iFlpMosaics* with scRNA-seq analysis is a very powerful tool for the high-throughput determination of cell-autonomous gene function in all cell types during development or disease.

**Figure 7:**
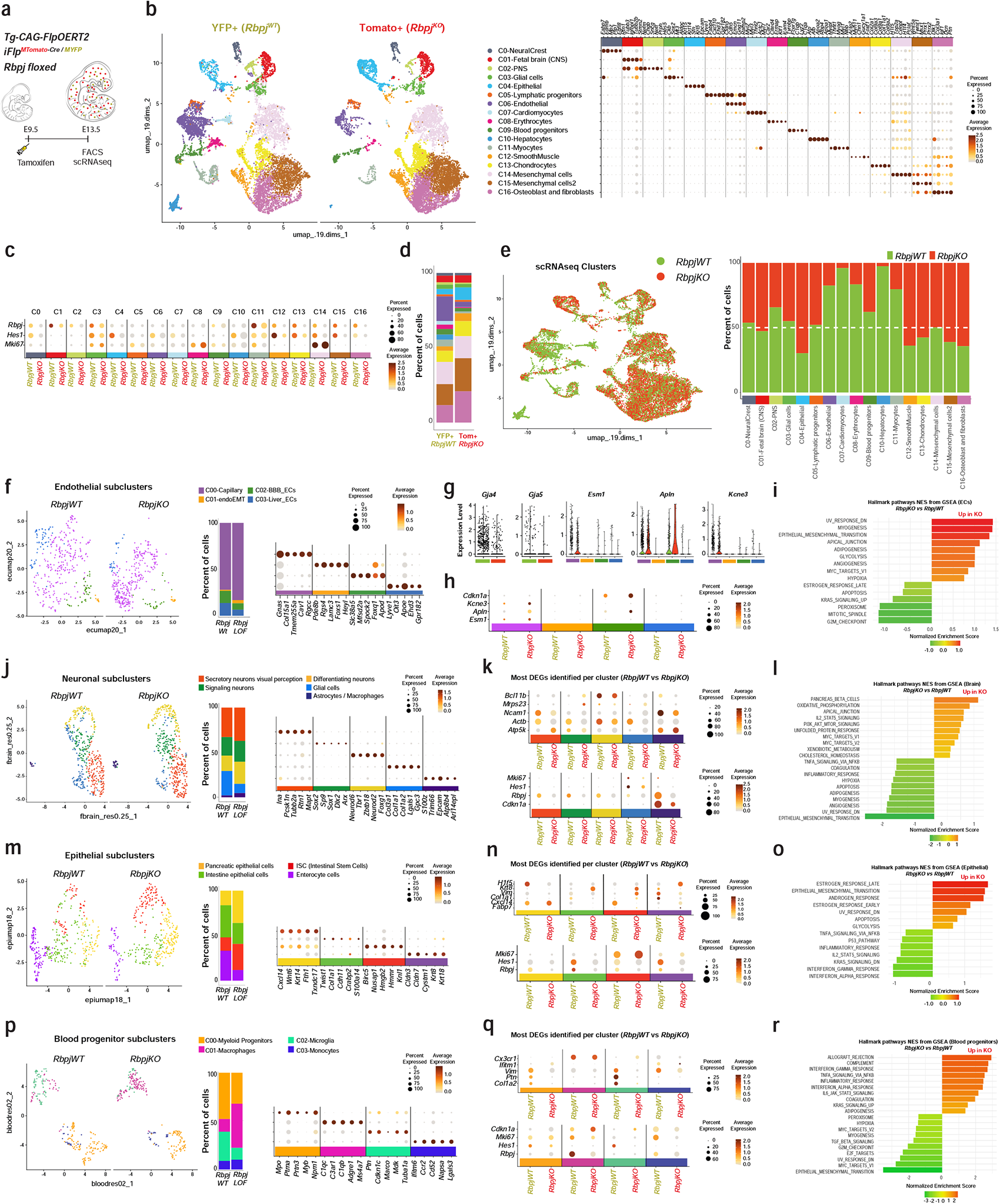
Combining single-cell RNA-seq with ratiometric *iFlpMosaics* to reveal cell-autonomous gene function. **a,** Schematic of the induction of the indicated alleles at E9.5 and collection of cells at E13.5 by FACS for scRNA-seq. **b**, Umaps showing the spatial distribution of the *Rbpj^KO^* (MTomato+) and *Rbpj^Wt^* (MYFP+) cells and the different clusters they form. The dot plot on the right shows the top markers used to identify the major cell types. **c**, Dot plot showing the frequency and amplitude of expression for *Rbpj,* the canonical Notch target *Hes1*, and the G2/M phase (proliferation) marker *MKi67* in wildtype cells (MYFP+) and mutant cells (MTomato-2A-Cre+). Note that *Rbpj* deletion only removes exons 6 and 7, leaving the 3’mRNA intact. However, this truncated mRNA is less stable and less detectable after deletion. **d**,**e**, Histobars and Umap showing that mosaic deletion of *Rbpj* (MTomato+ cells) changes the relative proportion of different cell types. **f-r**, Umaps showing the cell-type specific subclusters, together with top dysregulated marker genes and gene-set-enrichment analysis charts, showing how *Rbpj* deletion changes the frequency, differentiation, and biology of the indicated cell types.

## Discussion

Genetic mosaics are essential tools for determining the cell-autonomous function of a gene and avoid the confounding effects frequently associated with classical whole-animal or tissue-specific conditional genetics. With inducible mosaic genetics, the mutant and wildtype cells share the same microenvironment and developmental or physiological history, with the only difference between them being the deleted gene. This provides a much more accurate way to study and understand cell-autonomous gene function during tissue development, homeostasis, or disease.

We and others have observed that the transcriptional programs and phenotypes of single mutant cells in genetic mosaics often differ significantly from those of single cells extracted from a tissue in which the same gene was deleted in most cells (Hansen *et al*., 2022). Comparative omics of mutant and wildtype cells in genetic mosaics will more accurately reveal the true molecular and cellular gene cell-autonomous function. With classical genetics, changes to the mutant tissue function and microenvironment can always impact the transcriptional program of the mutant cells in ways unrelated to the original gene function or genetic program. For example, if a gene mutation impairs vascular development or permeability, this will affect the growth, function, and paracrine signaling of the surrounding tissue, and these effects will in turn influence the mutant-cell phenotype. With inducible genetic mosaics these effects are controlled for, because the wildtype and mutant cells share the same tissue history and microenvironment.

Nevertheless, there are several difficulties with the methods most frequently used for mosaic conditional genetics in mice, the most widely used animal model in biomedical research. Genetic mosaics induced by CreERT2 are easy to achieve, but are unreliable (Fernandez-Chacon *et al*., 2019; Liu *et al*., 2013; Schmidt-Supprian and Rajewsky, 2007). These models also lack the co-induction of wildtype-cell labeling required for direct phenotypic comparison with mutant and wildtype cells in the same tissue. MADM is genetically very reliable, and provides an internal wildtype cell control, but is a technically cumbersome method that provides no temporal resolution in the induction and genetic analysis (Contreras *et al*., 2021; Zong, 2014; Zong *et al*., 2005).

The set of new genetic tools we describe here significantly improves on both of these older technologies. Unlike Cre-ERT2 genetics, *iFlpMosaics* provide clearly distinguishable and reliable fluorescent labels for both true mutant cells (Tomato) and true wildtype cells (YFP). The simultaneous and ratiometric induction of fluorescent labels in mutant and wildtype cells in the same tissue is an important feature that allows accurate measurement and comparison of the proliferation, differentiation, migration, and survival of these populations over time. We have shown that this ratiometric analysis is robust over short and long periods of time. The fluorescent markers are easily detectable and quantifiable by microscopic imaging or FACS. Unlike MADM, *iFlpMosaics* provide a high degree of inducibility at any specific time point and in any cell type, including quiescent cells present in adult mice, which will be very important for modeling the etiology of diseases caused by sporadic somatic mutations in quiescent cells, or the consequences of CRISPR-based gene targeting and editing, which are by their nature also mosaic. In addition, *iFlpMosaics* are compatible with all existing floxed mouse alleles, enabling mosaic deletion of one or multiple floxed genes in cis, which is essential to perform functional genetic epistasis with single cell resolution. With *iFlpMosaics,* instead of only 1 cell barcode (as in standard CreERT2 or MADM), up to 12 different multispectral barcodes are possible for mutant cells and up to 7 for wildtype cells. This significantly higher spectral barcode diversity and dual labeling (membrane and nuclei) further increases single-cell and clonal quantitative resolution during a pulse– chase genetic experiment.

These new genetic tools allowed us to significantly extend and refine previous findings on the roles of several genes (*Myc*, *Foxo1/3/4* and *Rbpj*) in the development and differentiation of most embryonic lineages, the single cell clonal expansion of postnatal tissues, or the competitive survival of cells during the normal homeostasis and turnover of adult tissues. The ability to induce ratiometric genetic mosaics in multiple cell types at the same time, and at both low and high frequencies, increases the ease and throughput of a gene cell-autonomous function analysis. With *iFlpMosaics*, determining a gene cell-autonomous function no longer requires the crossing of mice carrying the floxed gene with multiple independent CreERT2 lines (one for each cell type).

In *iFlpMosaics,* cells expressing MTomato also express Cre, a recombinase that is toxic when expressed at high levels (Loonstra et al., 2001; Naiche and Papaioannou, 2007; Rashbrook et al., 2022). However, we found that the expression of the *iFlpMosaic* alleles is relatively low in comparison with previously published Cre-expressing lines (Fernandez-Chacon *et al*., 2019). This could be due to the different DNA elements used and the presence of only one upstream insulator during Rosa26 targeting. The PGK-Neo cassette and the 2A peptide elements used in i*FlpMosaic* alleles have been shown to reduce expression or translation efficiency (Ema et al., 2006; Pontes-Quero *et al*., 2017; Sharma et al., 2012). The absence of Cre toxicity with *iFlpMosaics* was confirmed by the long-term analysis of the Tomato-2A-Cre+:YFP+ cell ratio *in vitro* and *in vivo*. This ratio remained constant over time on a wildtype genetic background, assuring the absence of deleterious effects caused by the permanent Cre expression, similar to what has been shown for hundreds of other existing Cre lines.

Single-cell biology during tissue development is highly heterogenous, and it is therefore important to be able to reliably induce and analyze genetic mosaics of mutant and wildtype cells derived from single progenitor cells and with a negligible level of false positives and false negatives. The established MADM technology is ideal for this (Contreras *et al*., 2021; Zong, 2014; Zong *et al*., 2005) but lacks temporal inducibility. With the new *iFlp/Dre^Progenitor^* allele, we were able to significantly increase the inducibility and precision of *iFlpMosaics*. The dual *iFlp/Dre^Progenitor^* allele is much more sensitive to both FlpOERT2/DreERT2 activity and results in a genetic cascade that includes a delay in the expression of FlpO. This delay is very convenient for generating different mutant and wildtype cells within the progeny of a single progenitor cell, something previously achievable only with MADM. The progeny cells will also permanently express the nuclear marker H2B-V5 and the surface marker hCD2, ideal for FACS or rapid magnetic cell isolation of both mutant and wildtype cells.

Establishing the biology of single-cell lineages with high detail and in a realiable and high-throughput manner will require integration of information derived from *in situ* single-cell lineage-tracing microscopy with information derived from new scOMICs and bioinformatic analysis methods. Our analysis shows how the combination of *iFlpMosaics* with FACS or scRNA-seq easily achieves high-throughput analysis of the cell-autonomous function of any gene in all cell types during development, homeostasis, or disease. Since mutant and wildtype cell mosaics can be induced in fewer than 5% of the animal’s cells, they will not severely affect tissue or organ development and homeostasis, allowing analysis of the transcriptional, epigenetic, proliferative, and differentiation state of all different types of mutant cells in a mostly wildtype environment. This means that determining cell-autonomous gene function no longer requires crosses of mice carrying the floxed gene with multiple independent CreERT2 lines (one for each cell type). The ubiquituous and strongly expressed FlpOERT2 and DreERT2 lines generated here allow induction of *iFlpMosaics* in all cell types at both low and high frequency respectively. This enables direct comparison of the molecular and cellular phenotypes of mutant and wildtype cells extracted from all tissues within the same embryo or animal.

*iFlpMosaics* will greatly facilitate the induction and high-throughput functional analysis of somatic mutations or genetic mosaics. This will be central to understanding how an individual or combinatorial set of genetic mutations affects the biology of single cells during tissue development, regeneration, or disease.

## Methods

### Mice

We generated and used Mus musculus lines on the C57BL6 or C57BL6×129SV, or B6CBAF1 genetic backgrounds. All mice were backcrossed to C57Bl6 for several generations. To generate mice for analysis, we intercrossed mice aged between 7 and 30 weeks. We do not anticipate any influence on our data of mouse sex. The following mouse lines were used and intercrossed: *Tg(Cdh5-CreERT2)* (Wang *et al*., 2010), *Tg(iSuRe-Cre)* (Fernandez-Chacon *et al*., 2019), *Notch1^flox/flox^* (Radtke et al., 1999), *Rbpj^flox/flox^* (Han et al., 2002), *Myc^flox/flox^* (de Alboran et al., 2001), *Mycn^flox/flox^*(Knoepfler et al., 2002), *Foxo1/3/4^flox/flox^* (Paik *et al*., 2007), *Rosa26-EYFP* (Srinivas et al., 2001), *Rosa26-iChr2-Mosaic* (Pontes-Quero *et al*., 2017), Tg-*iMb2-Mosaic* or *iCre^MYFP/Tomato/MTFP1^*(Pontes-Quero *et al*., 2017), *Gt(Rosa)26^tm3(CAG-FlpOERT2)Ali^*(Lao *et al*., 2012), *Gt(Rosa)26Sor^tm14(CAG-LSL-tdTomato)Hze^*(Madisen et al., 2010), and *Apln-FlpO* (Luo et al., 2021). The following mouse lines were produced in this study: *R26-iFlp^MTomato-Cre/MYFP^, Tg-iFlp^MTomato-H2B-GFP-Cre/MYFP-H2B-Cherry-FlpO^*, *Tg(INS-CAG-FlpOERT2), Tg(INS-CAG-DreERT2), Tg(Cdh5-FlpOERT2)*, *iFlp^Chromatin^-Mosaic*, and *Tg-iFlp/Dre^Progenitor^* .

The *Tg-iFlp^MTomato-H2B-GFP-Cre/MYFP-H2B-Cherry-FlpO^*, *Tg(INS-CAG-FlpOERT2)*, *TgiFlp^Chromatin^-Mosaic*, and *Tg-iFlp/Dre-Progenitor* mouse alleles were generated by standard DNA injection into mouse eggs and screening of several transgenic founders, as indicated in the main text or figures. The *R26*-*iFlp^MTomato-Cre/MYFP^* allele was generated by CRISPR-assisted gene targeting of the Rosa26 locus in G4 ES cells, using the guide GACTGGAGTTGCAGATCACGA_GGG (IdT DNA) and a donor DNA with the elements indicated in Figure 1. Targeted ES cells were used for mice production. The *Tg(Cdh5-FlpOERT2)* line was generated by CRISPR-Cas9-assisted gene targeting of the existing *Tg(Cdh5-CreERT2)* line (Wang et al., 2010) in mouse eggs, using the guide AAGCTTATCGATACCGTCGA_CGG and a donor DNA containing the Cdh5-FlpOERT2 sequence. *Tg(INS-CAG-DreERT2)* mice were generated by CRISPR-Cas9 using guide RNAs (GATGTCGAACTGGCTCATGG_TGG and AACAGGCGGATCTGCGTACG_CGG) targeting the existing FlpO sequence contained in the pre-screened *Tg(INS-CAG-FlpOERT2)* transgene and a donor DNA containing the *CAG-DreERT2* sequence. Injection mixtures consisted of 0.305 mM of the described crRNAs (IDT) and tracrRNA (IDT, catalog 1072533) and 20 ng/µL Cas9 nuclease (Alt-R® S.p. HiFi Cas9 Nuclease V3, 100 µg, Catalog 1081060). Founders were initially screened by PCR with the primers described in Supplementary Table 1.

To induce CreERT2, FlpOERT2, or DreERT2 activity in adult mice, 1g tamoxifen (Sigma-Aldrich, P5648_1G) was dissolved in 50 mL corn oil (stock tamoxifen concentration, 20 mg/mL), and aliquots were stored at -20C. Animals received intraperitoneal injections of 50 µL to 150 µL of this stock solution (total dose, 1-to-3 mg tamoxifen per animal at 40-to-120 mg/kg), as indicated in the figures.

For treatment of pregnant females, the tamoxifen was dissolved together with progesterone to reduce miscarriage (2mg tamoxifen and 1mg progesterone per mouse). To activate recombination in pups, 4-Hydroxytamoxifen (4-OHT) was injected at the indicated stages at a dose of 40 mg/kg or 4mg/kg body weight, as indicated in the figures. All mouse lines and primer sequences required for genotyping are provided in Supplementary Table 1.

All mouse husbandry and experimentation were conducted using protocols approved by local animal ethics committees and authorities (Comunidad Autónoma de Madrid and Universidad Autónoma de Madrid CAM-PROEX 177/14, CAM-PROEX 167/17, CAM-PROEX 164.8/20 and PROEX 293.1/22). The mouse colonies were maintained in racked individual ventilation cages according to current national legislation. Mice had dust and pathogen-free bedding and sufficient nesting and environmental enrichment material for the development of species-specific behavior. All mice had ad libitum access to food and water in environmental conditions of 45%–65% relative humidity, temperatures of 21– 24°C, and a 12 h/12 h light/dark cycle. To preserve animal welfare, mouse health was monitored with an animal health surveillance program that followed FELASA recommendations for specific pathogen-free facilities.

### Immunofluorescence on cryosections

For multispectral *iFlpMosaics* tissue harvesting, mice were euthanized in CO_2_ chambers, 10 mL of 50 mM KCl were injected into the left ventricles, and whole mice were perfused with 3.7%-4% formaldehyde (ITW Reagents, 252931), at pH 7. Explanted tissues were post-fixed for 2 h in 4% paraformaldehyde (PFA) (Thermoscientific, 043368.9M) in phosphate-buffered saline (PBS) at 4°C with gentle rotation. After three washes in PBS for 10 min each, organs were stored overnight in 30% sucrose (Sigma) in PBS. Organs were then embedded in OCT^TM^ (Sakura) and frozen at –80°C. Cryosections (35 µm) were cut on a cryostat (Leica), washed three times for 10 min each in PBS, and blocked and permeabilized in PBS containing 10% donkey serum (Millipore), 10% fetal bovine serum (FBS), and 1% Triton X-100. Primary antibodies were diluted in the same buffer and incubated with sections overnight at 4°C. This step was followed by three 10 min washes in PBS and incubation for 2 h with conjugated secondary antibodies (1:200, Jackson Laboratories) and DAPI in PBS at room temperature. After three washes in PBS, sections were mounted with Fluoromount-G (SouthernBiotech). All antibodies used are listed in Supplementary Table 2.

### Wholemount immunofluorescence of retinas

For postnatal mouse retina immunostaining, eyes were collected and fixed in ice-cold 4% PFA in PBS for 10 minutes. Eyes were then incubated n the same solution for a further 15 min at room temperature, washed once in PBS, and kept on ice. After microdissection with spring scissors (FST), retinas were fixed in 4% PFA for an additional 45 min at room temperature, followed by two PBS washes of 10 min each. Retinas were blocked and permeabilized for 1 h in PBTS buffer (0.3% Triton X-100, 3% FBS, and 3% donkey serum). Samples were then incubated overnight at 4°C with biotinylated isolectinB4 (Vector Labs, B-1205, diluted 1:50) and primary antibodies (Supplementary Table 2) diluted in PBTS buffer. After five washes (20 min each) in PBTS buffer diluted 1:2, samples were incubated for 2 h at room temperature with Alexa-conjugated secondary antibodies (Supplementary Table 2). After three washes of 30 min each in PBTS buffer (diluted 1:2) and two washes of 15 min each in PBS, retinas were mounted with Fluoromount-G (SouthernBiotech).

### In vivo EdU labeling and detection of EC proliferation

To detect EC proliferation in postnatal liver and pancreas, 20 μg/g body weight EdU (Invitrogen, A10044) was injected intraperitoneally into P1, P7, or P20 mice 4 h before dissection. Livers and pancreases were fixed in 4% PFA and processed for cryosectioning as described above. EdU signals were detected with the Click-it EdU Alexa Fluor 647 or 488 Imaging Kit (Invitrogen, C10340 or C10337). In brief, after all other primary and secondary antibody incubations, samples were washed according to the immunofluorescence staining procedure and then incubated with Click-iT EdU reaction cocktail for 40 min, followed by DAPI counterstaining.

### Image acquisition and analysis

For confocal scanning, immunostained organ sections or whole-mount retinas were imaged at high resolution with a SP8 Navigator confocal microscope fitted with a 10x, 20x, or 40x objective. Individual fields or tiles of large areas were acquired. All images shown are representative of the results obtained for each group and experiment. Animals were dissected and processed under exactly the same conditions. Comparisons of phenotypes or signal intensities were made using images obtained with the same laser excitation and confocal scanner detection settings. Fiji/ImageJ was used to threshold, select, and quantify objects in confocal micrographs. Manual and automatic ImageJ public plugins and customed Fiji macros were used for quantification.

### Fluorescence activated flow cytometry and sorting

Embryonic, postnatal, or adult tissues were dissociated before fluorescence activated flow cytometry or sorting (FC or FACS). Embryonic tissues were dissociated using the Miltenyi Biotec Tissue Dissociation Kit 2 (130-110-203). Postnatal and adult mouse tissues were first digested for 20min at 37°C with 2.5 mg/mL collagenase type I (Thermofisher), 2.5 mg/mL dispase II (Thermofisher), and 50 μg/mL DNAseI (Roche).

Dissociated samples were filtered through a 70-μm cell strainer, and cells were centrifuged (400g, 4°C for 5 min). Cell pellets were gently resuspended in blood lysis buffer (0.15 M NH_4_Cl, 0.01M KHCO_3_, and 0.01 M EDTA in distilled water) and incubated for 10 minutes on ice to remove erythroid cells. Cells were centrifuged (400g at, 4°C for 5 min), and cell pellets were gently resuspended in blocking solution (PBS without Ca^2+^ or Mg^2+^ and containing 3% dialyzed FBS (Termofisher)) and incubated at 4°C with shaking for 20min. Cells were centrifuged (300g at 4°C for 5 min), resuspended, and incubated for 30min at 4°C with APC rat-anti-mouse CD31 and APC-CY7 anti-CD45 (Supplementary table 2). Cells were then centrifuged (400g, 4°C for 5 min), resuspended, washed in PBS without Ca^2+^ or Mg^2+^, and centrifuged again, and cell pellets were resuspended in blocking solution. Cells were kept on ice until used for FC or FACS. DAPI (5 mg/mL) was added to the cells immediately before analysis. Cells were routinely analyzed with a LSRFortessa cell analyser or sorted in a FACS Aria Cell Sorter (BD Biosciences). BD FACS Diva V8.0.1 and Flow JO v10 were utilized for FACS data collection and analysis.

### Cell isolation for transcriptomic analysis

For quantitative real time PCR (qRT-PCR) analysis, approximately 20,000 DAPI-negative Tomato+ or YFP+ cells from dissociated tissues were sorted directly to RLT buffer (RNAeasy Micro kit – Qiagen), and RNA was extracted according to the manufacturer’s instructions and stored at –80°C. This RNA was later used for qRT-PCR or RNAseq analysis. For qRT-PCR, total RNA was retrotranscribed with the High Capacity cDNA Reverse Transcription Kit with RNase Inhibitor (Thermo fisher, 4368814). cDNA was preamplified with Taqman PreAmp Master Mix containing a Taqman Assay-based pre-amplification pool composed of a mix of the Taqman assays indicated in Supplementary Table 3. Preamplified cDNA was used for qRT-PCR using the same gene-specific Taqman Assays and Taqman Universal Master Mix in an AB7900 thermocycler (Applied Biosystems). Data were retrieved and analyzed with AB7900 software.

For RNA-seq analysis, embryos were dissociated using Miltenyi Biotec Tissue Dissociation Kit 2 (130-110-203). A 1.2 mL volume of dissociation solution was placed together with a single embryo in a 2 mL round bottom tube. Following a 5-min incubation in a 37°C water bath, tubes were placed inside a pre-warmed 50 mL falcon tube and incubated for 25 min on a MACSmix Tube Rotor. Cells were then resuspended with 30 strokes with a Gilson P1000 pipette, starting slow and increasing speed gradually. To the 1.2 mL cell suspension were added 3 mL of sorting buffer (10% FBS in Ca_2_-and Mg_2_-free PBS), and the resulting solution was transferred to a 5 mL syringe. This volume was pressed slowly through a small 70 µm strainer to remove clumps. Each embryo yielded around 30 million cells at this stage.

Cell suspensions were spun at 400g for 6 minutes, and pellets were detached and resuspended in 300 µL sorting buffer containing DAPI. DAPI-negative, single, and live Tomato+ and YFP+ cells were sorted to eppendorf tubes containing 300ul of sorting buffer. Cells (66k Tomato+ and 30k YFP+) were sorted using a 100 µm nozzle, 20 PSI and high purity scale and low flow rate (1) to preserve cell viability and decrease contamination. Sorted cells were spun at 500g for 5min and resuspended in 30-40 µL cell-capture buffer (Ca_2_-and Mg_2_-free PBS supplemented with 0.04% BSA). After counting cells in Countess 3 Automated Cell Counter (Thermo Fisher Scientific), two independent 10x genomics ports were loaded with either 16k Tomato+ cells (90% viability) or 16k YFP+ cells (88% viability).

### Next generation sequencing sample and library preparation

Next Generation Sequencing (NGS) experiments were performed in the CNIC Genomics Unit. For scRNA-seq experiments, single cells were encapsulated in emulsion droplets using the Chromium Controller instrument (10x Genomics). scRNA-seq libraries were prepared according to the manufacturer’s instructions. The aimed target cell recovery for each port was 10000 cells. Generated libraries were sequenced in a HiSeq4000 or NextSeq2000 system (Illumina).

### Transcriptomic data analysis

Transcriptomic data were analyzed by the CNIC Bioinformatics Unit, Alvaro Regano, and Irene Garcia.

The following pipeline was followed for scRNA-seq data processing and *in silico* cell-type selection. For alignment and quantification of gene expression, the reference transcriptome was built using mouse genome GRCm39 and Ensembl gene build v109 (https://ftp.ensembl.org/pub/release-109/fasta/mus_musculus). The WPRE-sv40pa sequences expressed in the Tomato+ and YFP+ samples were added to the reference. Gene metadata were obtained from the corresponding Ensembl BioMart archive. Reads from transcripts were processed, aligned, and quantified using the Cell Ranger v6.1.2 pipeline. Single-cell analysis was based on the Seurat (Hao et al., 2021) and DoubletFinder R packages. Low-quality cells were filtered out using the following criteria: total counts, >1000 and <55,000; genes detected, >500 and <7500; mitochondrial transcripts content, <15%; hemoglobin transcripts, <1%; ribosomal transcripts, <35. Counts were log-normalized and scaled, followed by principal component analysis (PCA) and clustering using the shared nearest-neighbors algorithm and Louvain clustering (settings as defaults except for the 2000 most variable genes, 24 principal components, and a resolution of 0.35). Clusters and cells were classified based on the SingleR method (Aran et al., 2019) using Blueprint ENCODE, the Human Primary Cell Atlas cell-type profile collection, and a scRNASeq mouse dataset from the celldex package. This identification was used to predefine the different cell types present in the dataset for the analysis. Doublets were removed using DoubletFinder, using a first annotation with 24 PCs, 0.25 pN, and a pK of 0.05 and the second annotation with 10 PCs, pN0.25, and pK 0.05. Cells classified as Singlets were preserved. Singlets were then reclustered using 19 PCAs and a clustering resolution of 0.35. Manual curation of the identified clusters was applied to confirm and finetune the identity of the clusters based on the scRNAseq bibliography.

### Copy number assay and qRT-PCR

To determine the copy number of *FlpO*-or *Cre*-expressing cassettes after transgenesis, genomic DNA was extracted from mouse tail biopsies. Briefly, blood samples were digested overnight in 500 μL proteinase K solution (10 mM Tris-HCl [pH 8.0], 100 mM NaCl, 10 mM EDTA [pH 8.0], 0.5% sodium dodecyl sulfate, 0.25 mg/mL proteinase K) with occasional vortexing. To this solution were added 250 μL each of phenol and chloroform, followed by vigorous vortexing to ensure thorough mixing. Samples were immediately microcentrifuged at maximum speed for 5 minutes to separate the aqueous and organic layers. The upper aqueous layer (300 μL) was removed, with special care taken not to disturb the interface. To 300μl solution were added 30μL (0.1 x volume) of 3M NaAc and 825 μL (2.5 x volume) of 100% EtOH. The tubes were then shaken 10 times to precipitate the DNA. Samples were spun down at maximum speed for at least 45 minutes at 4°C. The supernatant was removed and pelleted DNA was washed with 500 μL 70% ethanol, followed by centrifuging at max speed for 5 minutes. The washed pellets were resuspended overnight in 100 μL TE solution (Tris 10mM and 0.1mM EDTA, pH 8). The next day, DNA concentrations were measured in a NanoDrop microvolume spectrophotometer (Thermo Fisher Scientific) and diluted to a final concentration of 10 ng/μL.

For the copy number assays, we performed qRT-PCR with TaqMan Universal Master Mix (TaqMan, 4440049) and the following probes: Tert TaqMan Copy Number Reference Assay (TaqMan probe, 4458369); a probe and primer set designed in the lab and synthetized by iDT to detect *FlpO* (qPCR FlpO probe: TCTTGATGTCGCTGAACCTGCCG [fluorophore FAM], FlpO-qPCR-F: CTGTACCAGTTCCTGTTCCTG, FlpO-qPCR-R: CTTGTCTTGGTCTCGGTCAC); and a pre-designed Cre assay to detect the *Cre* gene (Mr00635245_cn, ThermoFisher).

The relative number of copies of the genes of interest was determined using the Tert probe as a reference. Tert and FlpO/Cre probes were combined in the same reaction since they emit different fluorophores, FAM and VIC. qRT-PCR reactions were run in an AB7900 thermocycler (Applied Biosystems).

### Statistical analysis and reproducibility

All bar graphs show mean ± s.d. Experiments were repeated with independent animals. Comparisons between two sample groups with a Gaussian distribution were by unpaired two-tailed Student *t*-test. Comparisons among more than two groups were by one-way ANOVA followed by multiple comparison tests. All calculations were done in Excel, and final datapoints were analyzed and plotted with GraphPad Prism. No randomization or blinding was used, and animals or tissues were selected for analysis based on their genotype, the detected Cre/FlpO/Dre-dependent recombination frequency, and the quality of multiplex immunostaining. Sample sizes were chosen according to the observed statistical variation and published protocols.

## Data availability

RNA-seq data can be viewed at the Gene Expression Omnibus (GEO) under accession number GSEXXXX. Instructions and code to reproduce all scRNA-seq results can be found at https://github.com/RuiBenedito/Benedito_Lab. Unprocessed original photos of the data are available upon request. All other data supporting the findings in this study are included in the main article and associated files.

## Acknowledgments

Research in the Benedito laboratory was supported by European Research Council (ERC) Starting Grant AngioGenesHD (638028) and Consolidator Grant AngioUnrestUHD (101001814), the Ministerio de Ciencia e Innovación (SAF2017-89299-P and PID2020-120252RB-I00), and “la Caixa” Banking Foundation (project code HR19-00120 and HR22-00316 AngioHeart). The CNIC is supported by the Instituto de Salud Carlos III (ISCIII), the Ministerio de Ciencia e Innovación (MCIN), and the Pro CNIC Foundation and is a Severo Ochoa Center of Excellence (grant CEX2020-001041-S funded by MICIN/AEI/10.13039/501100011033).

Microscopy experiments were performed in the CNIC Microscopy and Dynamic Image Unit, an ICTS-ReDib co-funded by MCIN (/AEI /10.13039/501100011033) and the EDRF “A way to build Europe” (#ICTS-2018-04-CNIC-16).

I.G.-G. was supported by a PhD fellowship from Fundación La Caixa (CX-SO-16-1). A.R. was supported by The Youth Employment Initiative (YEI) PEJD-2019-PRE/BMD-16990. L.G.-O. was supported by the Spanish Ministry of Economy and Competitiveness (PRE2018-085283). S. G. was supported by a Juan de la Cierva Fellowship (FJC2020-044237-I). M.F.-C. was supported by PhD fellowships from Fundación La Caixa (CX_E-2015-01).

We thank S. Bartlett (CNIC) for English editing and the members of the CNIC Transgenesis, Microscopy, Genomics, Citometry, and Bioinformatic units. We also thank F. Alt (Boston Children’s Hospital – Harvard Medical School), T. Honjo (Kyoto University Institute for Advanced Studies), F. Radtke (Swiss Institute for Experimental Cancer Research), R. H. Adams (Max Planck Institute for Molecular Biomedicine), and R. De Pinho (MD Anderson Cancer Center) for sharing the *Myc ^floxed^, Rbpj ^floxed^, Notch1^floxed^, Cdh5(PAC)-creER, and Foxos1/3/4^floxed^* mice, respectively. We also thank Master’s students Joan Marc Arbonés and Alejandro Rivera Sánchez for their work on *iFlpMosaics*.

## Author contributions statement

I.G-G. and R.B. designed most of the experiments, interpreted results, assembled figures, and wrote the manuscript. R.B. designed all DNA vectors. I.G-G. did most of the DNA engineering (cloning), CRISPR-Cas9 genome targeting, animal experiments, confocal microscopy, and FACS analysis. Mice were generated by the CNIC Transgenesis Unit. S.G. developed new methods for *iFlpMosaics* induction, tissue immunostaining, and multispectral microscopy imaging, performed animal experiments and image analysis, and interpreted results. L.G-O, W.L., I.Z., and M.F-C. performed animal experiments, FACS, histology, and confocal imaging. I.G-G and A.R analyzed scRNA-seq data.

A.R. cloned and validated the *iFlp/Dre^Progenitor^* allele. S.F.R. performed image quantifications in Fiji and graphpad, edited text and figures, and assembled figures. M.L, M.S.S-M, A.G.-C, F.L., and V.C-G gave general technical assistance with experiments and genotyped the mouse colonies. All authors approved the final version of the manuscript.

## Figure legends

**Supplementary Figure 1:**
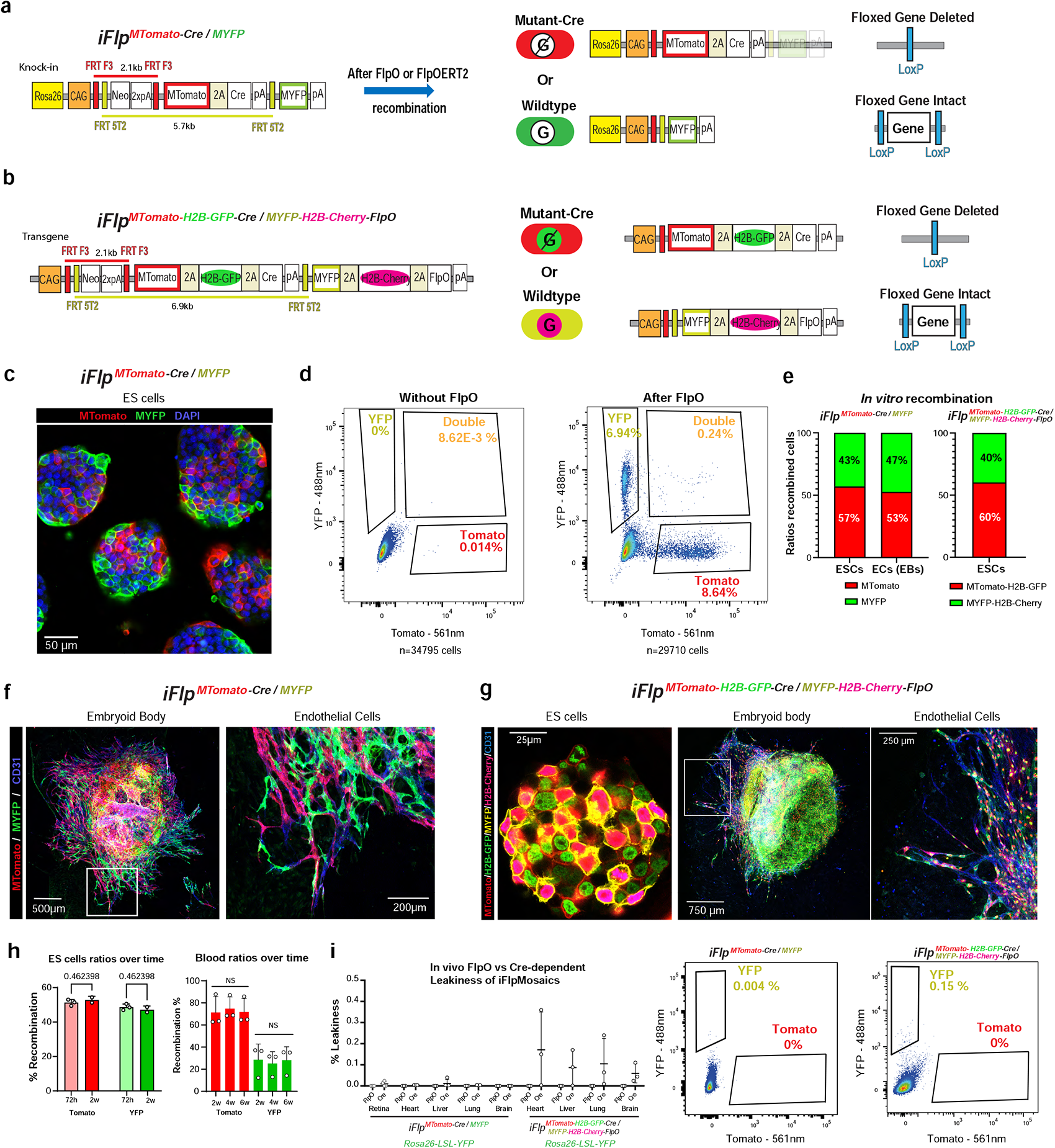
*iFlpMosaics* are neither toxic nor leaky. **a, b,** Schematic diagrams of the novel Rosa26-*iFlp^MTomato-Cre/MYFP^* and Tg-*iFlp^MTomato-H2B-GFP-Cre/MYFP-H2B-Cherry-FlpO^*alleles, showing the genetic distances between the mutually exclusive FRT site pairs and the expected outcomes after FlpO/FlpOERT2 recombination. **c, d** Confocal microscopy and FACS analysis of mouse ES cells used to generate Rosa26-*iFlp^MTomato-Cre/MYFP^* mice 3 days after transfection with FlpO-expressing plasmids. **e-g**, Frequency of recombination and expression detected by microscopy in ES cells and ECs derived from embryoid bodies (EBs). **h**, Relative frequency of MTomato-2A-Cre+ and MYFP+ cells in ES cells (in vitro) and in blood (in vivo); the absence of change over time shows that permanent expression of Cre is non-toxic to cells. **i**, FACS analysis of FlpO versus Cre-dependent leakiness in adult organs of mice carrying the standard *Rosa26-LSL-YFP* allele in combination with *iFlpMosaics* alleles. FlpO leakiness was not detected (0% Tomato+ cells), and Cre-non-self-leakiness (only detectable with the additional *Rosa26-LSL-YFP* allele) was observed only in a very small fraction of cells from animals carrying the Tg-*iFlp^MTomato-H2B-GFP-Cre/MYFP-H2B-Cherry-FlpO^* allele and not at all in animals carrying the Rosa26-*iFlp^MTomato-Cre/MYFP^*allele.

**Supplementary Figure 2:**
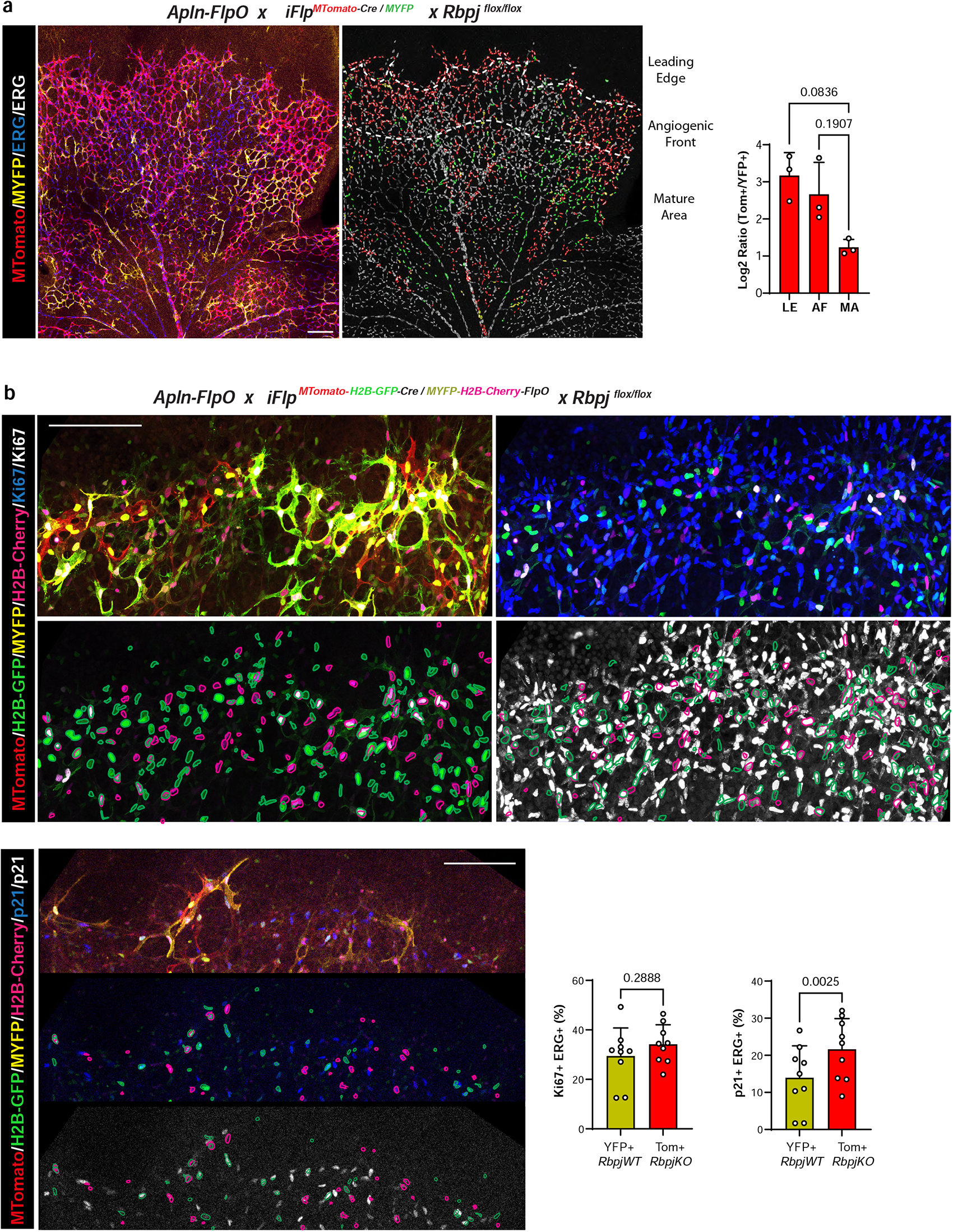
Validation of *iFlpMosaics* for ratiometric quantitative gene function analysis. **a**, Representative confocal micrographs of a P6 retina from a *Rbpj*-floxed mouse carrying the *iFlp^MTomato-Cre/MYFP^* and *Apln-FlpO* alleles. *Apln* is an X-chromosomal gene (mosaicly expressed in females) expressed at the angiogenic front of growing vessels, recombining in angiogenic front ECs that later form the entire vasculature. *Apln* also recombines in non-endothelial (ERG-negative) cells in the retina. The right micrograph shows ERG-signal segmentation (EC nuclei), with nuclei of Tomato+ (*Rbpj^KO^*) cells in red and nuclei of YFP+ (*Rbpj^WT^*) cells in green. Grey nuclei are in non-recombined ERG+ cells. Dashed white lines demark the leading edge (LE), angiogenic front (AF), and mature area (MA). A, arteries; V, veins. The chart shows the log2 ratios of Tomato+ to YFP+ ECs in each retinal region, showing the enrichment of *Rbpj^KO^* cells in the LE and AF, which suggests a role for this gene in cell migration to the front of the plexus. **b, c** Representative confocal micrographs of a P6 retina from *Rbpj*-floxed mouse carrying the *iFlp^MTomato-H2BGFP-Cre/MYFP-H2BCherry^*and *Apln-FlpO* alleles. This allows the nuclear labeling of *Rbpj^KO^*(H2B-GFP+) and *Rbpj^Wt^* (H2B-Cherry+) ECs and blood cells, which is more convenient for cell object segmentation and quantification of nuclear-specific proteins, such as the proliferation and cell-cycle arrest markers Ki67 and p21. Nuclear segmentation outlines: red, Cherry+/Ki67-or p21-; pink, Cherry+/Ki67+ or p21+; green, GFP+/Ki67-or p21-; cyan, GFP+/Ki67+ or p21+. Scale bars, 150μm.

**Supplementary Figure 3:**
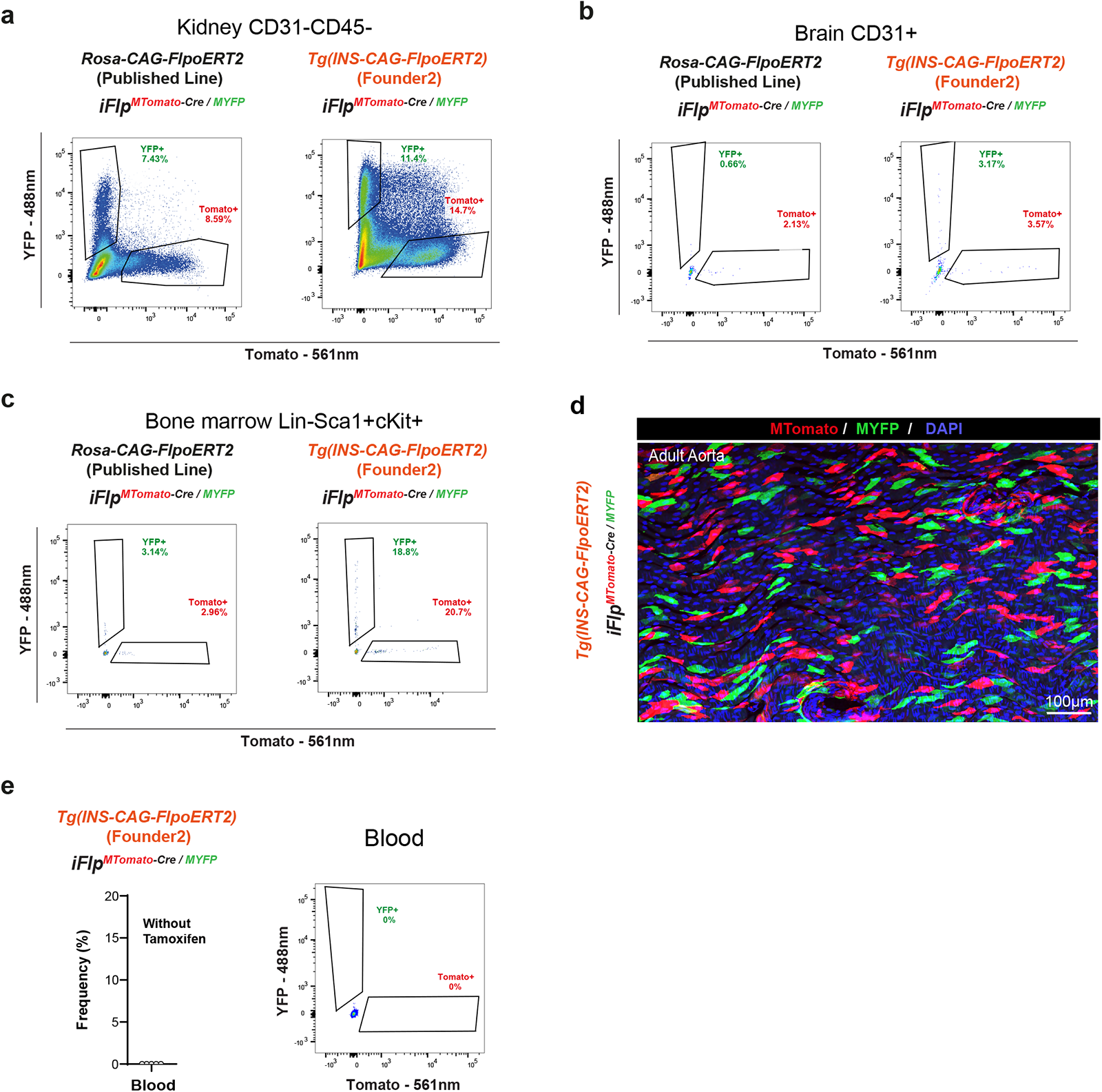
Comparative analysis of recombination efficiency in FlpOERT2 mouse lines. **a-c**, Representative FACs plots of different adult organs, showing YFP+ and Tomato+ cell frequencies in from mice carrying the *iFlp^MTomato-Cre/MYFP^*allele in combination with the published *R26^CAG-FlpoERT2^* or the newly generated *Tg(Ins-CAG-FlpoERT2*)^F2^ allele. **d,** Confocal micrograph of a flat-mounted adult mouse aorta 2 weeks after induction. **e,** FACS analysis showing the lack of leakiness of the *Tg(Ins-CAG-FlpoERT2*)^F2^ allele. In the absence of tamoxifen, the *iFlp^MTomato-Cre/MYFP^* allele is not recombined.

**Supplementary Figure 4:**
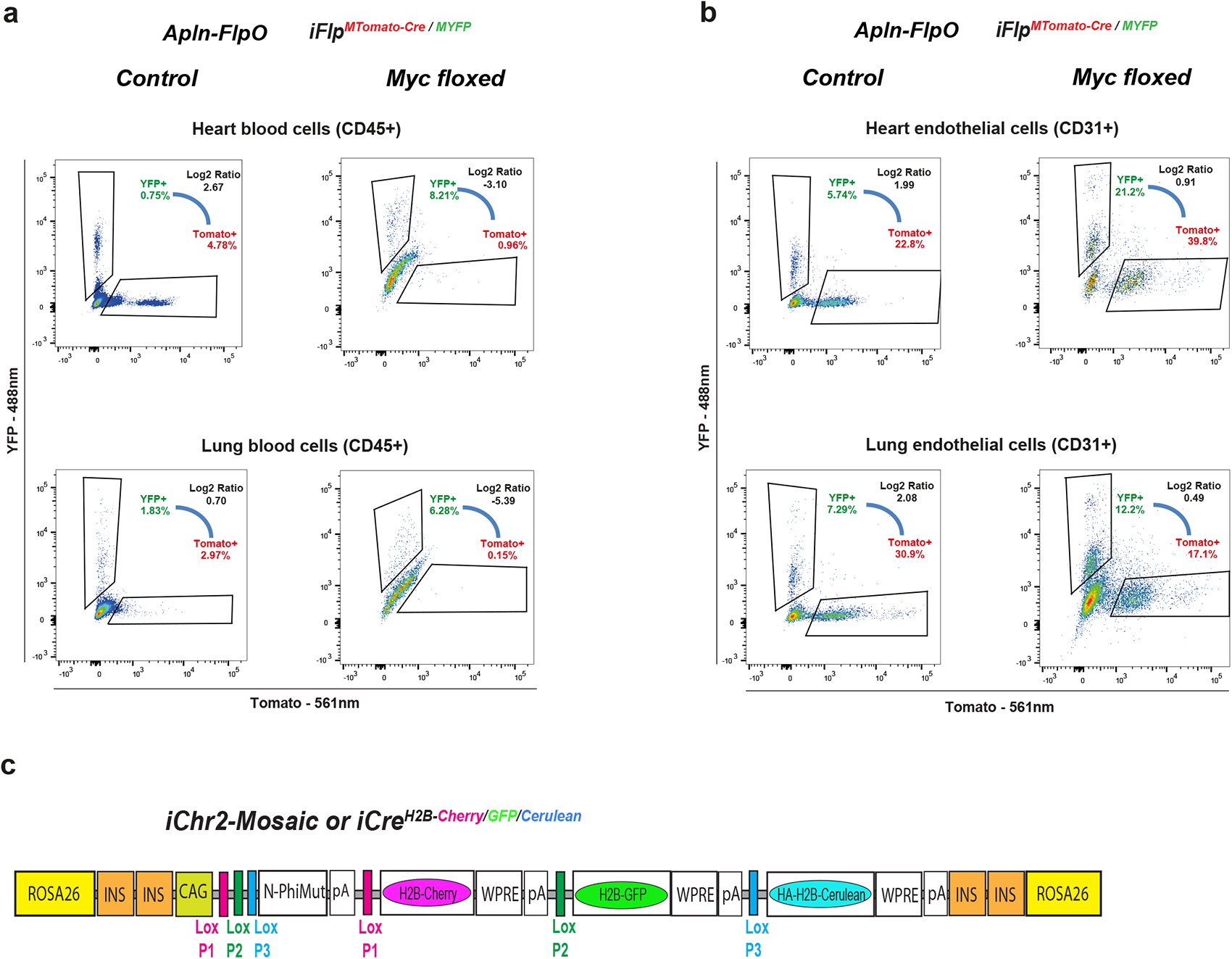
Ratiometric iFlpMosaics reveal that the competitive disadvantage of *Myc^KO^*cells is stronger in blood cells than in ECs. **a,b**, Representative FACS plots showing that in both the heart and the lung CD45+ blood cells are much more affected by *Myc* deletion than CD31+ ECs. c, Schematic diagram of the *iChr2-Mosaic* construct (also called *iCre^H2B-Cherry/GFP/Cerulean^*) targeted to the Rosa26 locus and having 4 insulator (INS) sequences and the strong and ubiquituous CAG enhancer/promoter. N-PhiMut encodes a non-fluorescent nuclear marker protein detectable with an antibody. It is expressed before Cre recombination. After Cre-mediated recombination between compatible pairs of LoxP sites, only one of the 3 possible fluorescent proteins will be expressed. WPRE, woodchuck hepatitis virus post-transcriptional regulatory element, used to enhance transgene expression. pA, Sv40 polyA, used to terminate transcription.

**Supplementary Figure 5:**
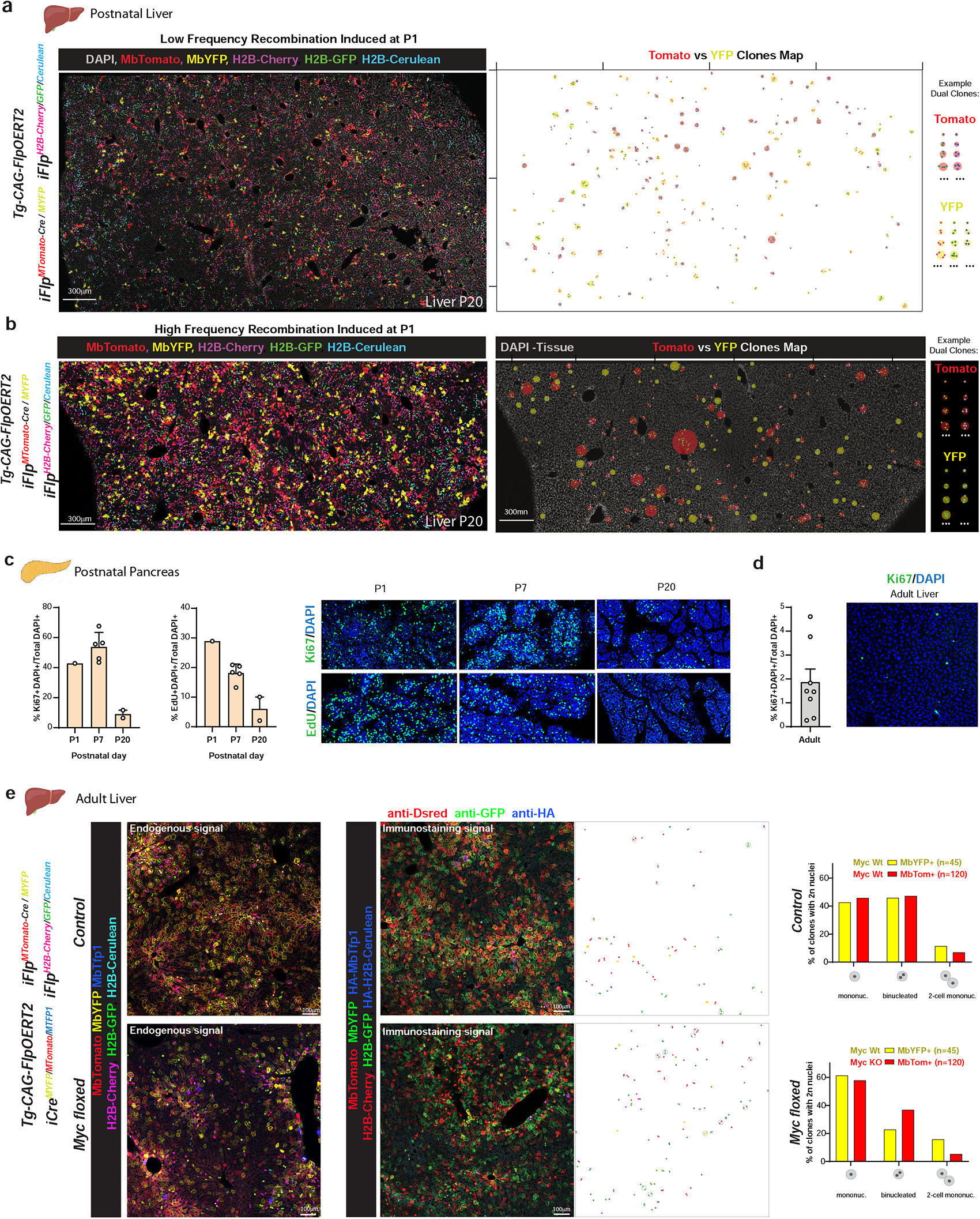
Mapping the clonal expansion of single mutant and wildtype cells with *iFlpMosaics*. **a,b**, Representative confocal images (low magnification large tile scans) of entire P20 liver sections from mice carrying the indicated alleles. Recombination of the reporter alleles was induced at P1 and resulted in different recombination frequencies. Right panels show the results of automatic image processing with Fiji scripts, which generate spatial maps of dual-labeled clones of cells (with membrane and nuclear labeling). The bigger the circle, the bigger the clone. **c,** Quantification and representative confocal micrographs of proliferative pancreatic cells (Ki67+ in cycle or EdU+ in s-phase) at different postnatal stages. **d,** Quantification and representative confocal micrograph of Ki67+/DAPI+ cells in the adult liver. **e,** Representative confocal micrographs of adult liver sections from control and *Myc*-floxed mice carrying the indicated alleles. Left panels show endogenous fluorescent signals from the reporter alleles. Center panels show sections from the same mice immunostained with antibodies to DsRed (detects MTomato and H2B-Cherry), GFP (detects MYFP and H2B-GFP), or the HA epitope (detects MTFP1 and H2B-Cerulean) together with Fiji-script-generated spatial maps of dual-labeled clones (with membrane and nuclear labeling) reveal hepatocyte ploidy and clone size. Graphs show the percentages of mononucleated cells, binucleated cells, and 2-cell mononuclear clones. In control mice, MYFP+ (WT) and MTomato+ (WT) diploid cells occur in similar proportions, whereas in *Myc*-floxed mutants MTomato+ (*Myc^KO^*) diploid cells are more frequently binucleated and give rise to fewer 2-cell mononucleated clones than MYFP+ (*Myc^WT^*) cells.

**Supplementary Figure 6:**
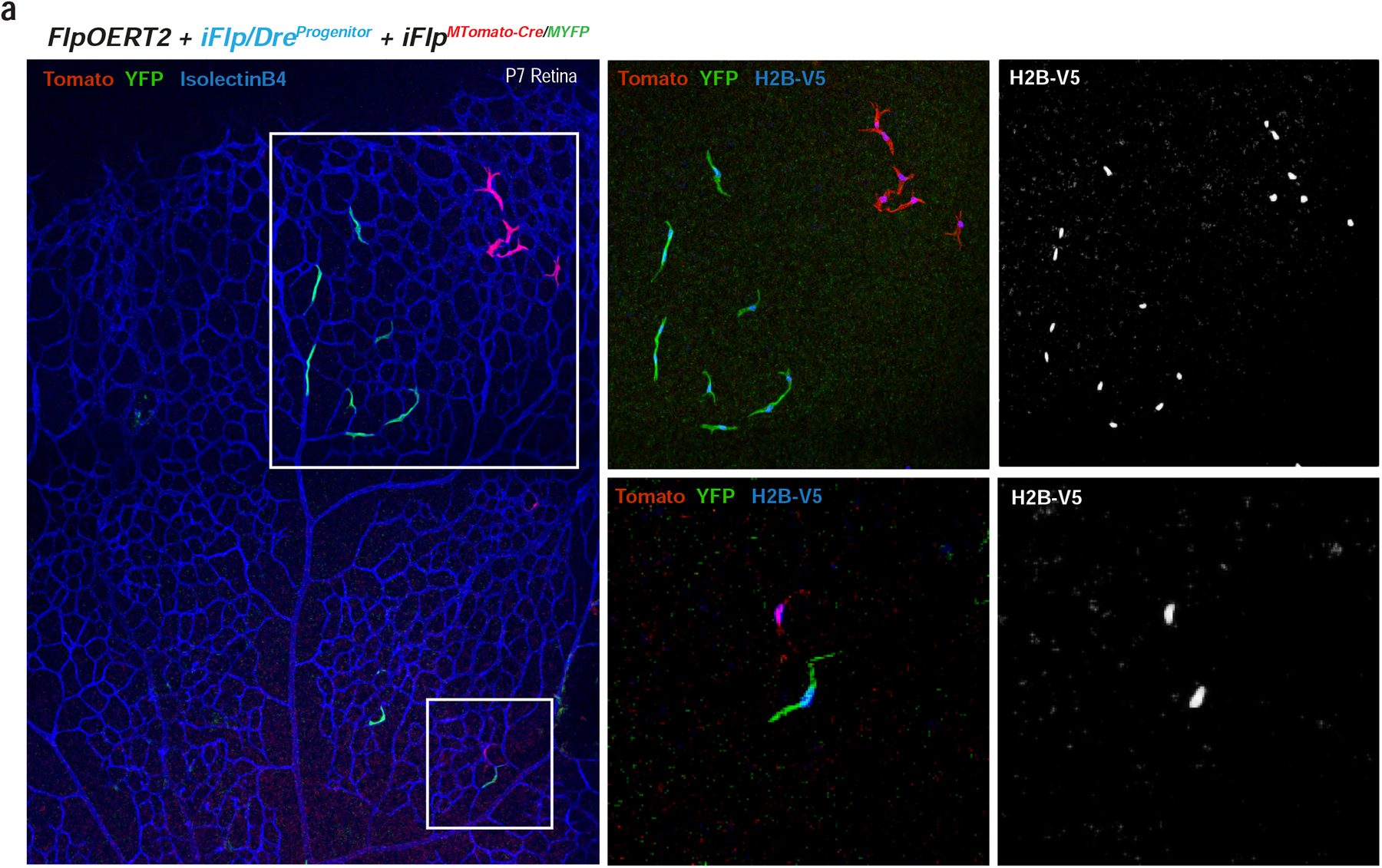
*iFlp/Dre^Progenitor^*enables the induction of genetic mosaics from a single progenitor cell. **a**, Representative confocal micrograph of a P7 retina from an animal carrying the indicated alleles and induced with 4-OHT at P3. After induction of the ***iFlp/Dre^Progenitor^***allele, cells express the nuclear marker H2B-V5 and FlpO. After some rounds of cell division, FlpO will recombine the *iFlp^MTomato-Cre/MYFP^* allele, generating MYFP+ and MTomato+ progeny cells derived from a single initial recombination event.

**Supplementary Figure 7:**
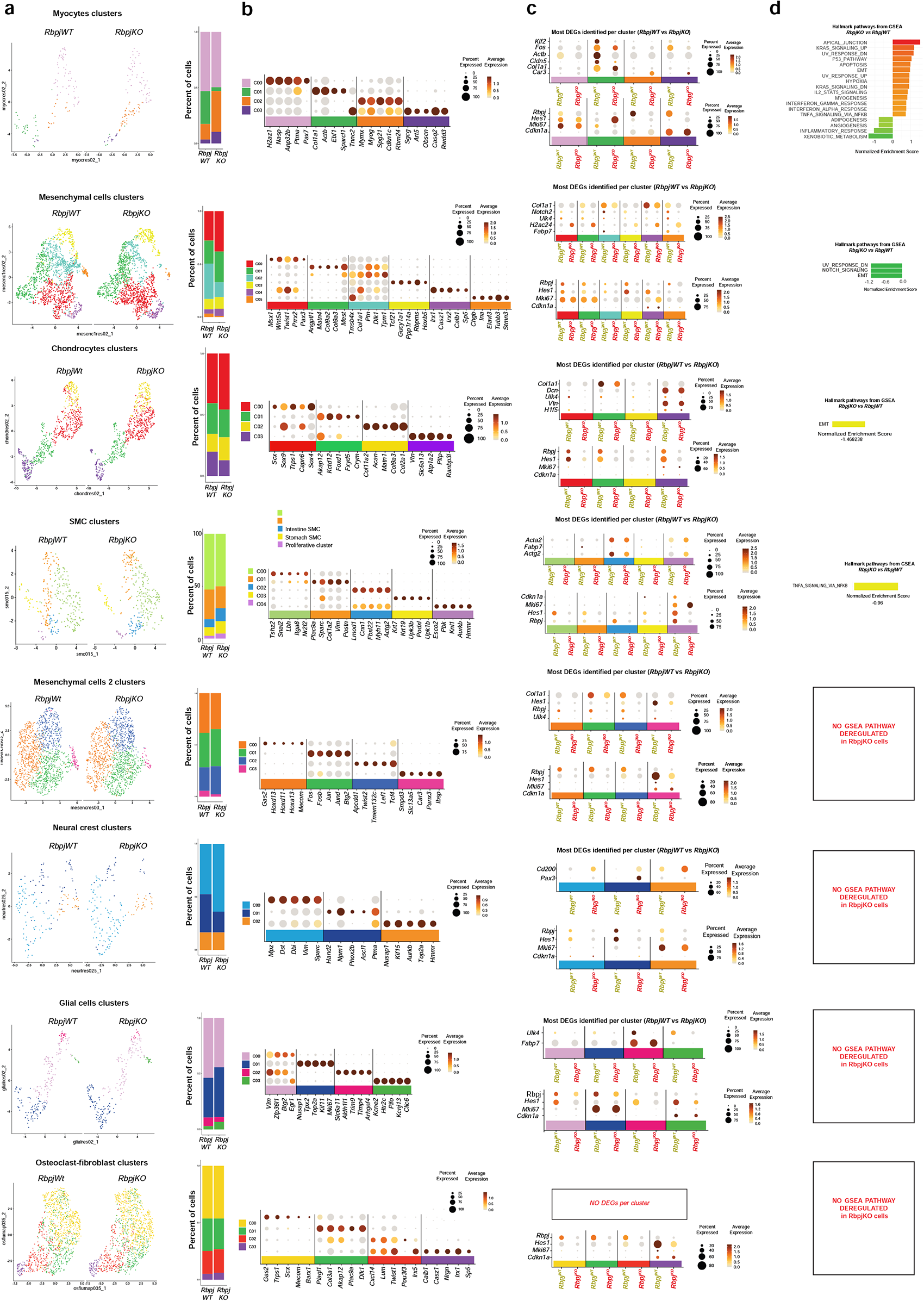
Single cell RNA-seq analysis combined with ratiometric iFlpMosaics uncovers cell-autonomous gene function in diverse cell types. **a,** Umaps and barplots showing the identified clusters and their frequencies among YFP+ (*Rbpj^WT^*) and Tomato+ (*Rbpj^KO^*) cells. **b,** Dot plots showing the frequency (size) and expression level (color intensity) for the top cluster marker genes. **c,** Top differentially expressed genes per cluster between YFP+ (*Rbpj^WT^*) and Tomato+ (*Rbpj^KO^*) cells. Most of the charts also show the expression of *Rbpj*, its canonical target *Hes1*, and the proliferation markers *Ki67* (G2/M cells) and *Cdkn1a* (arrested cells). **d,** Gene set enrichment analysis (GSEA) pathways with their normalized enrichment score. For some cell types there are no differentially expressed genes or pathways.

**Supplementary Table 1.**
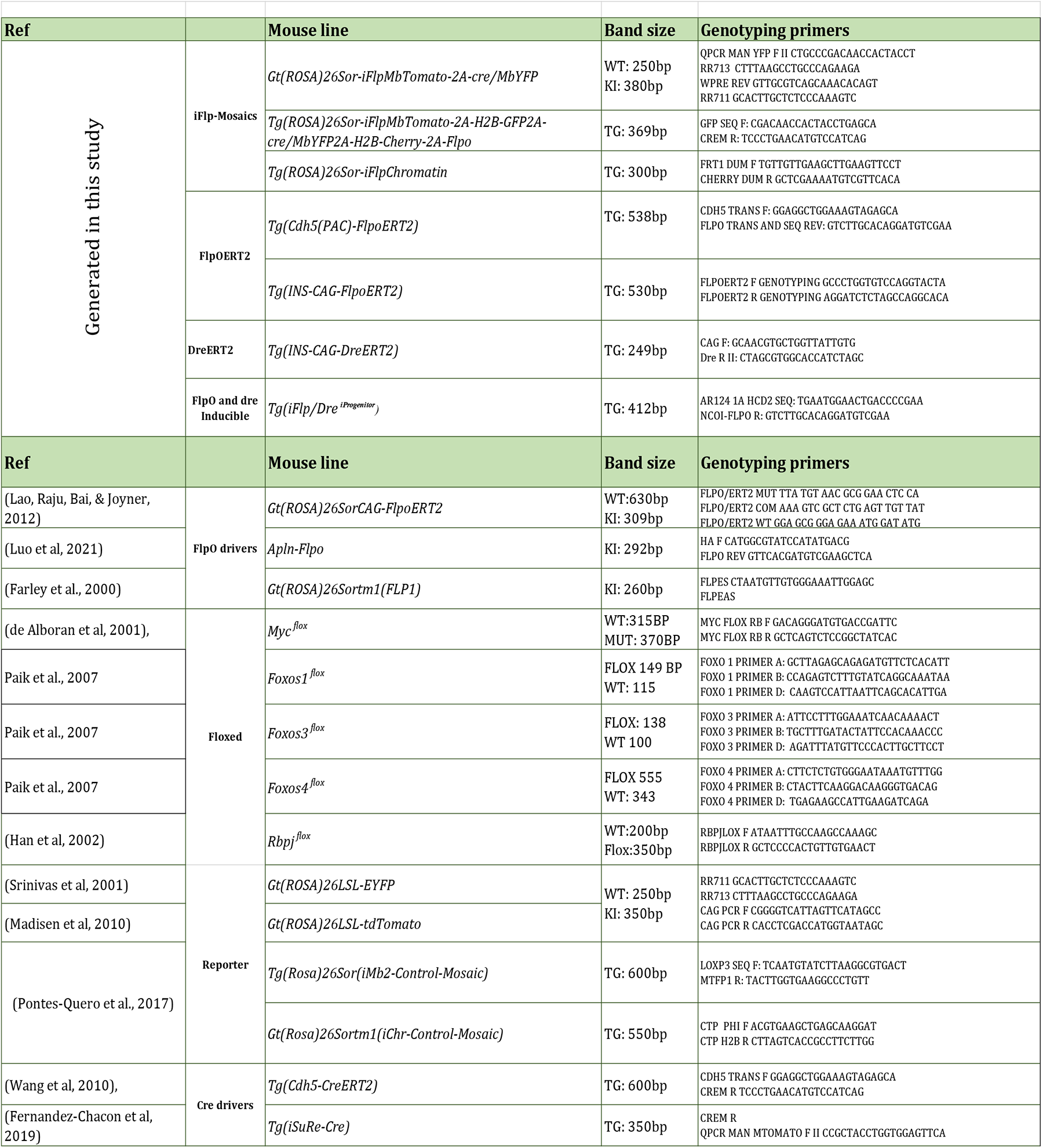

**Supplementary Table 2.**

| Antibody | Dilution | Source | Reference |
| --- | --- | --- | --- |
| **Primary Antibodies** |  |  |  |
| Biotinilated Isolectin B4 | 1:100 (IF) | VECTOR LABS | B-1205 |
| Rat anti- mouse p21 Rat monoclonal Clone: HUG0291 Supernatant | 1:50 (IF) | CNIO | # |
| Monoclonal Rabbit anti-ERG [EPR3863] | 1:500 (IF) | ABCAM | Ab110639 |
| Goat Polyclonal Anti-GFP and YFP/CFP variants | 1:200 (IF) | Acris | R1091P |
| Anti-PECAM-1 Antibody, clone 2H8 hamster anti mouse CD31 | 1:200 (IF) | Merck | MAB1398Z |
| Rabbit anti- DsRed (Living colors) | 1:200 (IF) | Clontech | 632496 |
| Rabbit anti-RFP/Dsred/Tomato/Cherry CF543 conjugated (Polyclonal) | 1:200 (IF) | Biotium | 20476 |
| Mouse anti HA-Tag (6E2) Mouse mAb (Alexa Fluor® 647 Conjugate) | 1:200 (IF) | Cell Signaling | 3444 |
| RABBIT ANTI-RFP/Dsred ConjugatedCF594 | 1:200 (IF & FC) | Sigma or Biotium | BTIU20422 |
| Rabbit Anti-ERG Alexa Fluor® 647 EPR3864 | 1:500 (IF) | ABCAM | ab196149 |
| Rat anti-Ki-67 Monoclonal Antibody (SolA15), eFluor 660, eBioscience™ | 1:250 (IF) | Thermo Fisher Scientific | 50-5698-82 |
| APC Rat Anti-Mouse CD31 | 1:250 (FC) | BD Bioscience | 551262 |
| APC-Cyanine7 Rat Anti-Mouse CD45.2 | 1:250 (FC) | TonboBio | 25-0454-U025 |
| Mouse anti HA-Tag (6E2) Mouse mAb (Alexa Fluor® 647 Conjugate) | 1:250 (IF) | Cell Signaling | 3444 |
| BV711 Rat Anti-Mouse CD31 Clone MEC 13.3 (RUO) | 1:250 (FC) | BD Biosciences | 740680 |
| Rabbit anti GFP Conj. Alexa-488 | 1:200 (IF & FC) | Invitrogen | A213111 |
| BD Horizon™ BV605 Rat Anti-Mouse CD45 | 1:200 (FC) | BD Biosciences | 563053 |
| Anti-Mouse CD150 (SLAM) Antibody Brilliant Violet 510™ | 1:300 (FC) | Biolegend | 115929 |
| Anti-Mouse CD48 APC Cy7 | 1:100 (FC) | Biolegend | 103432 |
| Anti-Mouse CD117 (c-kit) PercP Cy5.5 | 1:200 (FC) | Biolegend | 105824 |
| Anti-Mouse Ly-6A/E (Sca-1) FITC | 1:200 (FC) | eBioscience | 11-5981-85 |
| Biotin Mouse Lineage Depletion Cocktail | 1:200 (FC) | BD Biosciences | 51-9000794 |
| **Secondary Antibodies** |  |  |  |
| Streptavidin, Alexa Fluor® 405 conjugate | 1:200 (IF) | Thermo Fisher Scientific | ST32351 |
| Donkey Anti-Rat IgG H&L (Alexa Fluor® 647) preadsorbed | 1:400 (IF) | ABCAM | ab150155 |
| Cy™3 AffiniPure Fab Fragment Donkey Anti-Rabbit IgG (H+L) | 1:400 (IF) | Jackson | 711-167-003 |
| AffiniPure Goat Anti-Armenian Hamster Alexa680 | 1:400 (IF) | Jackson | 127-625-16 |

**Supplementary Table 3.** Taqman Assays

| Gene Symbol | Dye | Assay ID | Source |
| --- | --- | --- | --- |
| *Rbpj* | FAM-MGB | Mm01217627_g1 | Thermo Fisher |
| *Myc* | FAM-MGB | Mm00487804_m1 | Thermo Fisher |
| *Mycn* | FAM-MGB | Mm00476449_m1 | Thermo Fisher |
| *Notch1* | FAM-MGB | Mm00435245_m1 | Thermo Fisher |
| *Pecam1* | FAM-MGB | Mm01242584_m1 | Thermo Fisher |
| *Gapdh* | FAM-MGB | Mm99999915_g1 | Thermo Fisher |

